# Systematic multi-level analysis of an organelle proteome reveals new peroxisomal functions

**DOI:** 10.1101/2021.12.08.471723

**Authors:** Eden Yifrach, Duncan Holbrook-Smith, Jérôme Bürgi, Alaa Othman, Miriam Eisenstein, Carlo W.T Van Roermund, Wouter Visser, Asa Tirosh, Chen Bibi, Shahar Galor, Uri Weil, Amir Fadel, Yoav Peleg, Hans R Waterham, Ronald J A Wanders, Matthias Wilmanns, Nicola Zamboni, Maya Schuldiner, Einat Zalckvar

**Author notes:** co-correspondence.

## Abstract

Seventy years following the discovery of peroxisomes, their proteome remains undefined. Uncovering the complete peroxisomal proteome, the peroxi-ome, is crucial for understanding peroxisomal activities and cellular metabolism. We used high-content microscopy to uncover the peroxi-ome of the model eukaryote – *Saccharomyces cerevisiae*. This strategy enabled us to expand the known organellar proteome by ∼40% and paved the way for performing systematic, whole-organellar proteome assays. Coupled with targeted experiments this allowed us to discover new peroxisomal functions. By characterizing the sub-organellar localization and protein targeting dependencies into the organelle, we unveiled non-canonical targeting routes. Metabolomic analysis of the peroxi-ome revealed the role of several newly-identified resident enzymes. Importantly, we found a regulatory role of peroxisomes during gluconeogenesis, which is fundamental for understanding cellular metabolism. With the current recognition that peroxisomes play a crucial part in organismal physiology, our approach lays the foundation for deep characterization of peroxisome function in health and disease.

## Introduction

All eukaryotic cells, from yeast to humans, compartmentalize various functions into membrane-enclosed organelles. One such organelle, the peroxisome, is crucial for human health and survival as it hosts essential metabolic enzymes (Waterham et al., 2016). Despite its importance to organismal health and cellular metabolism, the full repertoire of proteins that reside within peroxisomes had not been fully uncovered.

One substantial leap that fueled the scientific community’s interest in other metabolic organelles such as the mitochondria was the formation of its protein compendium, the MitoCarta (Pagliarini et al., 2008). To put peroxisomes into the limelight, and expose the complete variety of peroxisomal functions, we set out to uncover the peroxisomal proteome (the peroxi-ome).

The discovery of the entire peroxi-ome is important not only in the diagnosis and treatment of patients suffering from peroxisomal diseases but much more broadly in the study of viral infection and immunity (Cook et al., 2019), malignant transformation (Kim, 2020), aging (Titorenko and Terlecky, 2011; Pascual-Ahuir et al., 2017) and neurodegenerative diseases (Zarrouk et al., 2020) – all of which have now clearly demonstrated a role for peroxisomes in their progression.

However, discovering the complete peroxi-ome is a challenging task – peroxisomes are small and physically attach to multiple other organelles via contact sites (Schrader et al., 2020; Shai et al., 2016; Valm et al., 2017; Castro et al., 2018). Peroxisomes also change dramatically in response to cell state or environmental conditions (Smith and Aitchison, 2013).

Various systematic studies were previously used to discover peroxisomal proteins. Most efforts relied on either mass-spectrometry based analysis of peptides in subcellular fractions, or on sequence-based strategies to detect proteins with a potential canonical Peroxisomal Targeting Signal 1 or 2 (PTS1 or PTS2) (Schrader and Fahimi, 2008), which are known to be recognized by peroxisomal targeting factors (Walter and Erdmann, 2019). However, both of these approaches have limitations such as the difficulty in detecting low-abundance and conditionally expressed proteins by subcellular fractionations, or the inability to find non-canonically targeted proteins by sequence-based approaches. Based on these efforts, as well as low-throughput protein-specific studies, we have recently curated comprehensive lists of peroxisome proteomes in Humans, Mice (Yifrach et al., 2018), and in *Saccharomyces cerevisiae* (from herein called yeast) (Yifrach et al., 2016). These efforts highlighted that proteins responsible for known peroxisomal activities (Grunau et al., 2009; Antonenkov and Hiltunen, 2012) have not yet been described and suggested that new approaches are necessary to identify more peroxisomal proteins.

We, therefore, decided to take a complementary approach and sought to map the peroxi-ome by performing a high-content screen on fluorescently tagged yeast proteins. Here we report the identification of 33 new peroxisomal proteins, which together with the known ones make the most complete inventory of the peroxi-ome to date, containing 115 proteins in total (Data S1). Having a comprehensive view of the peroxi-ome enabled us to create a new strategy to characterize peroxisomal activities – one that relies on a systematic analysis of the entire peroxi-ome at various functional levels. Alongside targeted assays, we uncovered several uncharted peroxisomal functions. For example, by systematically studying the sub-organellar localization and targeting mode of the entire organellar proteome, we exposed non-canonical targeting dependencies on the main targeting factor, Pex5. By proteome-wide analysis of the metabolomic profiles of peroxisomal mutants, we revealed unexpected metabolic functions in peroxisomes. Importantly, we identified a novel mechanism by which peroxisomes regulate gluconeogenesis, a process that generates sugars from non-carbohydrate substrates. This finding exposes an unexpected link between peroxisomal activity and gluconeogenesis that goes beyond the provision of building blocks following fatty acid degradation. More broadly, the identification of tens of new peroxisomal proteins paves the way for additional exciting discoveries regarding peroxisome function and regulation and introduces a more holistic view on this important, yet understudied organelle.

## Results

### A high-content screen maps the peroxi-ome

To uncover the peroxi-ome we performed a high-throughput microscopic screen on a recently-made full-genome yeast collection harboring a Green Fluorescent Protein (GFP) tag fused to the amino terminus (N’) of each yeast protein (Weill et al., 2018). This strategy has several advantages: (i) The cells are analyzed by imaging keeping all cellular structures intact. This is especially important for proper identification of proteins that are localized to several compartments and could be regarded as contaminants in mass-spectrometry based analysis of enriched peroxisomal fractions as they are not enriched in these fractions, (ii) The method is unbiased and does not rely on any prior knowledge on the protein sequence, (iii) Expression is regulated under a constitutive *NOP1* promoter, which enables us to detect the localization of low-abundance and conditionally expressed proteins, (iv) Having a GFP tag on the N’ of each protein allows proper targeting of all known proteins with either a PTS1 or a PTS2 motif (Yofe et al., 2016) and (v) We have previously performed a pilot experiment and verified that this methodology works well in identifying new peroxisomal proteins (Yifrach et al., 2016).

To unequivocally identify peroxisomal structures we integrated a peroxisomal marker, Pex3-mCherry, to the entire yeast library using an automated mating procedure (Cohen and Schuldiner, 2011; Tong and Boone, 2006) and screened the new, custom-made library, containing both the N’ GFP tagged proteins and the peroxisomal marker using an automated fluorescence microscope (Fig. 1A). To sensitize the screen, we increased the number and size of peroxisomes by growing yeast on the fatty acids oleate as a sole carbon source, a condition that makes peroxisomes larger, more numerous, and essential for yeast survival. Moreover, some proteins, although expressed in glucose, are targeted to peroxisomes only in oleate-containing media, as we have previously shown (Yifrach et al., 2016). To ensure that we are not missing proteins that co-localize to peroxisomes only in glucose we performed an additional screen in glucose-containing medium only for 280 strains that were annotated to have a punctate localization in glucose in the original N’ GFP library (Weill et al., 2018).

**Figure 1.**
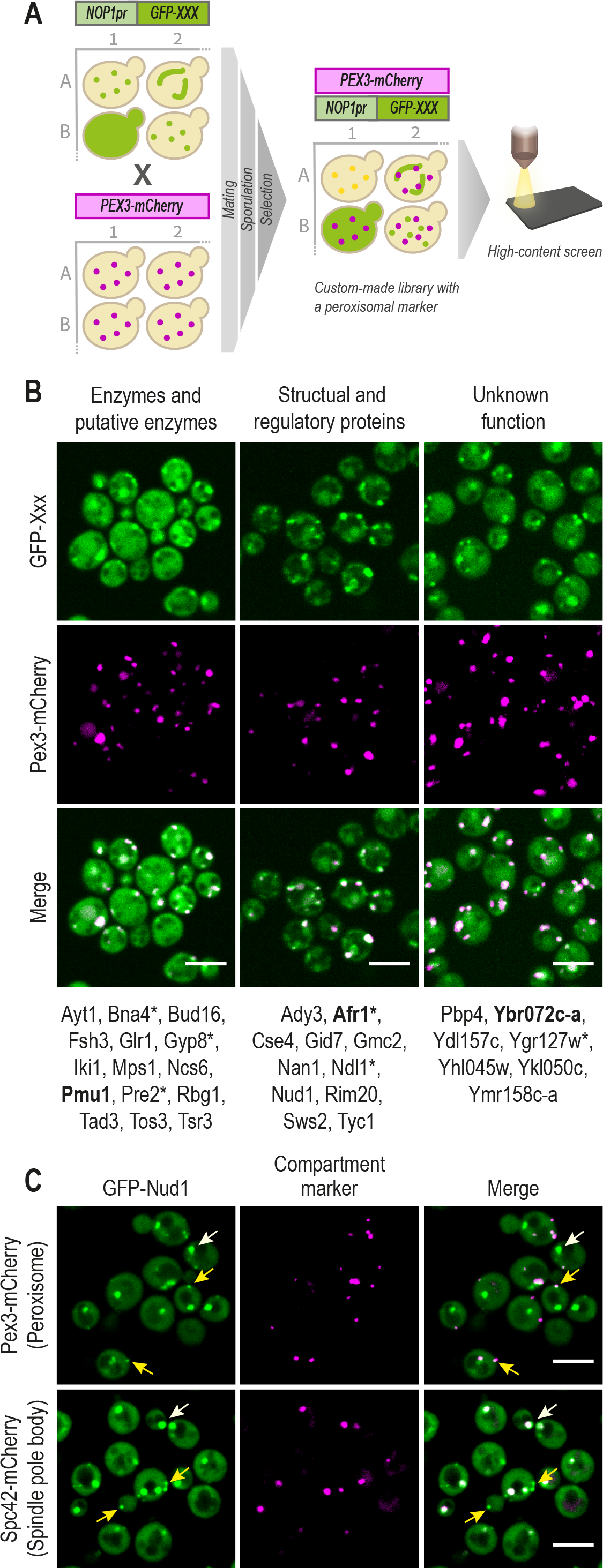
A high-content screen maps the peroxi-ome. (A) A peroxisomal marker Pex3-mCherry was genomically integrated into a yeast N′ GFP collection by utilizing an automated mating procedure. Then, high-throughput fluorescence microscopy was applied to identify proteins that co-localize with the peroxisomal marker. Only proteins that passed three validation steps (manual retagging, PCR, and western blot (Fig. S1)) were determined as newly-identified peroxisomal proteins. (B) The validated proteins were divided into groups to represent their variety of functional annotations: enzymatic, structural, or unknown activity. In bold are the proteins whose images are presented. Asterisks mark proteins that were also detected in peroxisomes when their encoding genes are expressed under their native promoter. (D) Protein localization image analysis demonstrated that about half of the newly-identified proteins are dually-localized to peroxisomes as well as to other compartments. Presented is GFP-Nud1 that co-localizes with both a spindle-pole body marker (white arrows) and with a peroxisomal marker (yellow arrows). Full analysis of dual-localization is shown in Fig. S2 and Table S1. For all micrographs, a single focal plane is shown. The scale bar is 5 µm.

Following manual analysis of all images, we found 50 N’ GFP tagged proteins that co-localized with the peroxisomal marker and were not previously reported as peroxisomal proteins. We performed several verification steps including manual re-tagging with GFP of all 50 proteins, ascertaining correct genomic integration using PCR, re-imaging, and western blot analysis (Fig. S1). This stringent filtering narrowed down the initial list to 33 newly-identified peroxisomal proteins (hits) (Data S1). The newly-identified proteins expand the present protein count of peroxisomes by ∼40%. These proteins include enzymes and putative enzymes, structural and regulatory proteins as well as uncharacterized proteins of unknown function (Fig. 1B). We re-imaged all hits under their native promoter and found that all proteins that could be visualized above background when expressed from their native promoter were indeed peroxisomal (Fig. 1B, marked with asterisks), verifying that peroxisomal localization was not a result of the constitutive expression. The low expression levels of many of the proteins when expressed under the regulation of their own promoter may explain their absence in previous proteomics-based approaches.

Interestingly, ∼50% of the proteins in our list are dually localized to peroxisomes and other compartments such as mitochondria, nucleolus, and bud-neck (Fig. 1C and Fig. S2). This may explain why they were not previously identified as peroxisomal proteins and demonstrates the advantage of using an imaging-based approach. For example, Nud1 is a core component of the spindle pole body (SPB) outer plaque. We show that, as expected, GFP-Nud1 co-localizes with an SPB marker, but that it has an additional, secondary, localization to peroxisomes (Fig. 1C).

Overall, our approach uncovered tens of proteins that by fluorescence microscopy resolution seem to associate with peroxisomes, although they were not assigned as such before. This puts forward the most comprehensive peroxi-ome list, that can be now used to study different aspects of peroxisome biology using systematic, whole-proteome technologies, to discover new peroxisomal functions.

### High-resolution imaging reveals the sub-organellar distribution of the peroxi-ome

To further characterize the peroxi-ome, we wanted to uncover their sub-organellar distribution systematically. Regular fluorescence microscopes are limited to a resolution of ∼250 nm. This resolution is not sufficient for ascertaining co-localization between small cellular structures nor for detecting the sub-organellar distribution of proteins in most organelles. Therefore, we exploited a long-established observation in which deletion of the peroxisomal membrane protein Pex11 and incubation of cells in oleate-containing media causes to enlargement of peroxisomes (Erdmann and Blobel, 1995). We noticed that the C terminal (C’) tagging of Pex11 causes the same phenomena. Combining the genetic expansion with high-resolution microscopy provided us with a sufficient spatial resolution to differentiate between membrane- and matrix-localized organellar proteins.

Hence, we applied an automated mating procedure to integrate Pex11-mScarlet into a yeast collection where each strain expressed one protein of the peroxi-ome fused at its N’ to GFP and imaged all strains by high-resolution microscopy following growth in oleate-containing media (Fig. 2A).

**Figure 2.**
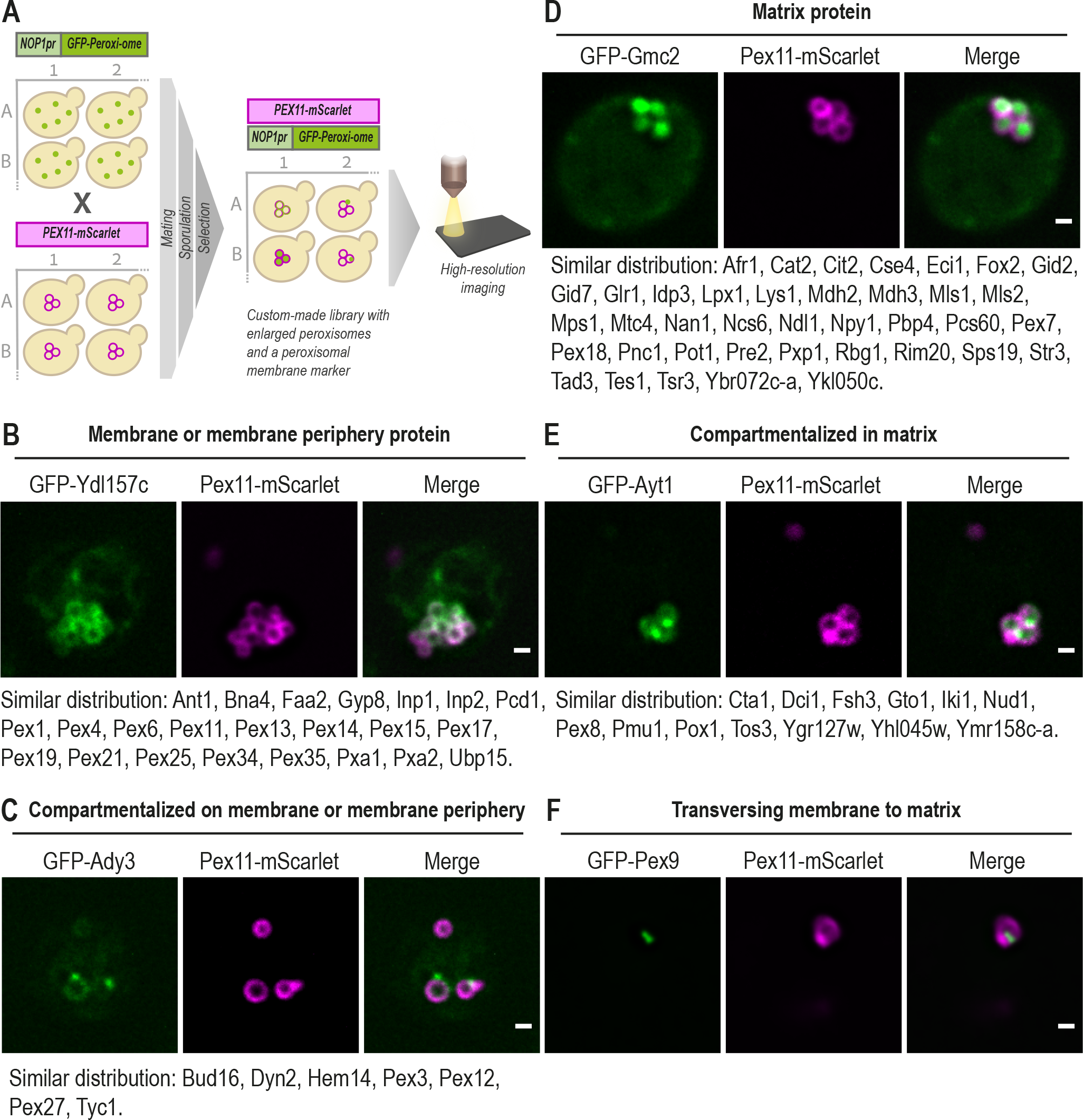
High-resolution imaging reveals the sub-organellar distribution of the peroxi-ome. (A) The sub-peroxisomal localization of each peroxi-ome protein was captured by high-resolution imaging and was enabled by genomic and metabolic enlargement of peroxisomes. Enlargement was induced by the integration of the C’ tagged form of the peroxisomal-membrane protein Pex11 into a yeast N′ GFP collection using an automated mating procedure and upon 20 hours incubation of cells in oleate-containing media (B) GFP-Ydl157c represents a protein that distributes homogeneously on the peroxisomal membrane or membrane periphery. (C) GFP-Ady3 represents a protein that compartmentalizes on the peroxisomal membrane or membrane periphery. (D) GFP-Gmc2 represents a protein that distributes homogeneously in the peroxisomal matrix. (E) GFP-Ayt1 represents a protein that compartmentalizes in the matrix. (F) GFP-Pex9 was the only protein that transversed the peroxisomal membrane. For each category, proteins with similar distributions are listed below the micrographs. For all micrographs, a single focal plane is shown. The scale bar is 500 nm.

Analysis of the entire peroxi-ome distribution revealed five different sub-peroxisomal localizations. (i) proteins that localize homogeneously on the peroxisomal membrane or membrane periphery, for example, Ydl157c (Fig. 2B), (ii) proteins that compartmentalize on the membrane or the membrane periphery, such as Ady3 (Fig. 2C), (iii) proteins that localize homogeneously in the peroxisomal matrix, like Gmc2 (Fig. 2D), (iv) proteins that compartmentalize within the matrix, such as Ayt1 (Fig. 2E) and (v) one protein, Pex9, that had a unique distribution that seems to transverse the membrane (Fig. 2F).

Our microscopic results (summarized in Data S1) go hand in hand with previous biochemical data curated for many known peroxisomal proteins and with trans-membrane domain (TMD) predictions made by systematic topology analysis within TopologYeast (Weill et al., 2019). Importantly, we reveal the distribution of all the newly-identified peroxisomal proteins and unveil the sub-organellar localization of several known peroxisomal proteins. For example, some proteins, such as Pex3, Pex12, Pex27, and Hem14, are not evenly distributed but rather seem to compartmentalize on specific sites of the membrane. While Pex3 was previously shown to compartmentalize on the membrane and form contact sites with the plasma membrane (Hulmes et al., 2020) and the vacuole (Wu et al., 2019), the other proteins were never suggested to have such a distribution. It will be interesting to further study whether this distribution is due to their function in membrane contact sites. Overall, exploring which membrane or matrix peroxisomal proteins compartmentalize with which may shed light on potential functional complexes and suggest new roles for peroxisomal proteins.

Furthermore, a unique sub-peroxisomal distribution was observed for Pex9, a conditional targeting factor that brings a subset of PTS1 proteins into the matrix (Yifrach et al., 2016; Effelsberg et al., 2016). Interestingly, Pex9 seemed to transverse the peroxisomal membrane (Fig. 2F). It was previously shown that each of the constitutive targeting factors, Pex5 and Pex7, form an import pore together with the docking protein Pex14 to insert folded, and even oligomerized, proteins through the peroxisomal membrane, into the matrix (Montilla-Martinez et al., 2015). Our microscopic results strengthen the recent finding that Pex9 requires the same set of peroxisomal membrane proteins to mediate the protein import (Rudowitz et al., 2020) and suggest that it can form a pore with the docking complex on the membrane to insert fully folded proteins into the organelle.

Overall, our sub-organellar analysis not only confirmed that all the newly-identified proteins are indeed intimately connected to peroxisomes, but it also provided a glimpse into the inner-peroxisomal protein organization and lays the foundations for building a more detailed map of peroxisome function.

### Functional mapping of targeting dependencies for matrix proteins reveals multiple non-canonical Pex5 substrates

Following our sub-organellar analysis, it was clear that most peroxisomal proteins are localized to the matrix, however, many of them do not contain a canonical targeting sequence that is recognized by one of the two main targeting factors, Pex5 and Pex7. Therefore, we wanted to systematically map the targeting pathways which these proteins utilize. To do this we visualized each of the GFP-tagged peroxisomal proteins on the background of either *PEX5* or *PEX7* deletions (Fig. 3A). Known Pex5 or Pex7 cargo proteins served as positive controls to ascertain that we can clearly distinguish specific cargos by this strategy (Fig. S3A). Interestingly, all newly-identified matrix proteins depended on *PEX5*, but not on *PEX7*, for their peroxisomal localization (Table S1 and example in Fig. S3B). Sequence analysis of the newly-identified matrix proteins showed that only six of them were previously predicted, yet not proven, to contain a PTS1 (Notzel et al., 2016), the targeting sequence that is known to be recognized by Pex5.

**Figure 3.**
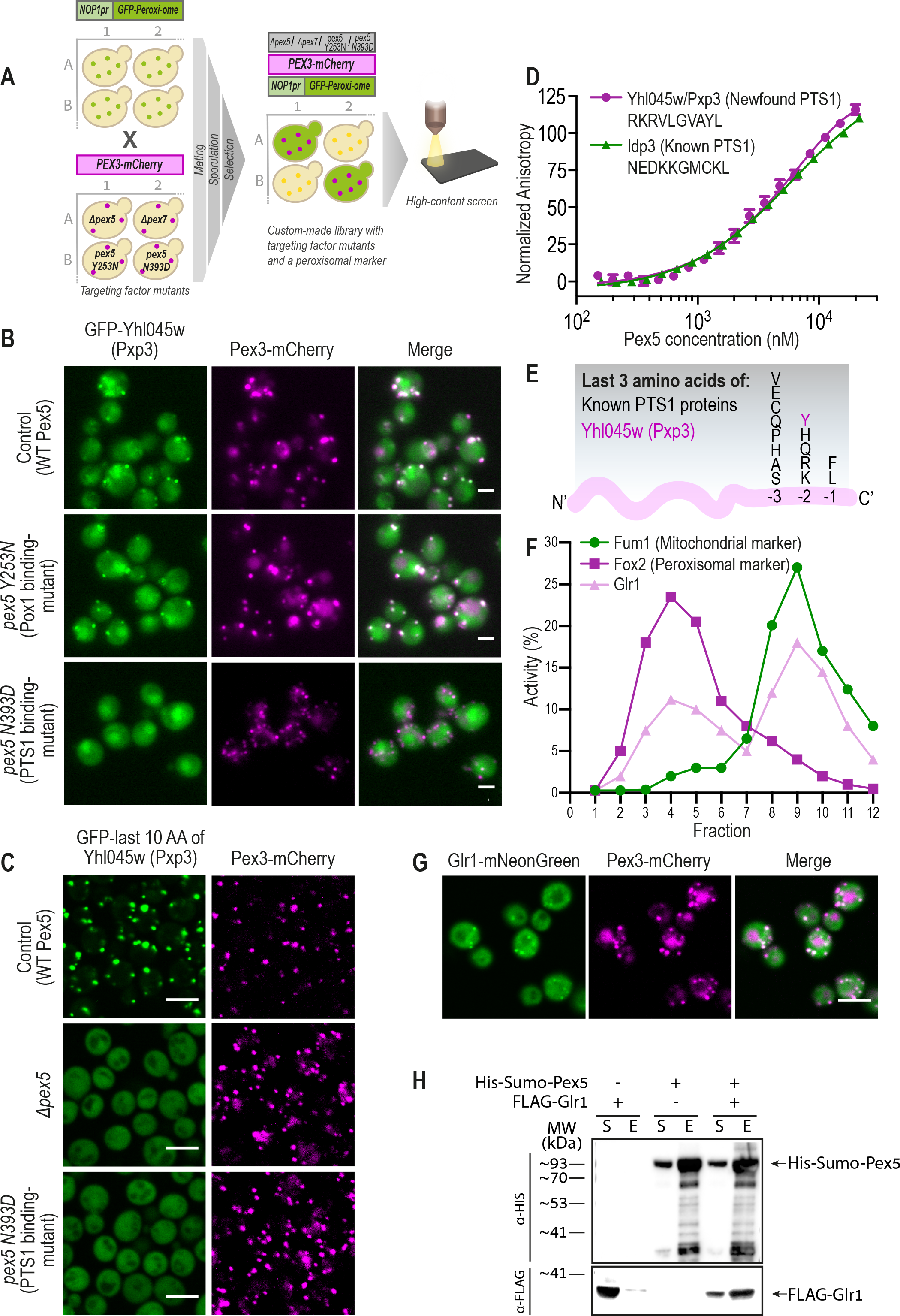
Functional mapping of targeting dependencies for matrix proteins reveals multiple non-canonical Pex5 substrates. (A) Targeting dependencies of the newly-identified peroxisomal matrix proteins were uncovered using a high-content screen. Targeting factor deletions (*Δpex5* or *Δpex7*) or point mutations (*PEX5*Y253N weakens the interaction with Pox1, a non-PTS1 protein, and *PEX5*N393D weakens the interaction with PTS1 proteins), were used to examine effects on peroxisomal localization of each newly-identified protein. A strain carrying a Hygromycin selection cassette in an inert locus was used as a control for no modification in peroxisomal genes. (B) Matrix proteins targeting was dependent on the PTS1 pathway of the Pex5 targeting factor (full analysis in Table S1). Presented is GFP-Yhl045w, which was not targeted to peroxisomes upon a point mutation in the PTS1-binding domain of Pex5 (*PEX5*N393D). (C) The peroxisomal targeting ability of the predicted motifs was examined by fusing the last 10 amino acids of each protein to the C’ of GFP, integrating the construct into an inert locus in the yeast genome, and imaging. While most motifs were unable to target GFP to peroxisomes (Fig. S4B), the Yhl045w (Pxp3) motif was sufficient to target GFP to peroxisomes, in a Pex5 PTS1-binding site-dependent manner. (D) A fluorescence anisotropy experiment demonstrated direct binding of the Pxp3 motif with purified Pex5 protein (Kd = 9.5 +/-2.9 µM), in strength similar to the binding of the known PTS1 motif from Idp3 (Kd = 5.5 +/- 0.5 µM). (E) The consensus sequence of PTS1 motifs in yeast is now extended with the newly-identified, unique, residue (Tyrosine, Y) at position −2 of Pxp3. (F) Subcellular fractionations followed by enzymatic activity assays show that untagged, native, Glr1 is active also in peroxisomes, in addition to its known activity in mitochondria. Fum1 and Pox2 were used as markers for mitochondrial, and peroxisomal fractions, respectively. (G) Glr1 C’ was fused to mNeonGreen and its peroxisomal localization was analyzed by fluorescence microscopy, demonstrating Glr1 does not rely on a free C’ for targeting to peroxisomes hence it does not contain a canonical PTS1 motif (H) *In vitro* pull-down assays analyzed by western blot show that Glr1 was co-eluted with Pex5, despite not having a canonical PTS1 motif. S-Soluble fraction, E-Elution fraction. For all micrographs, a single focal plane is shown. The scale bar is 5 µm.

How are the 22 proteins that depend on *PEX5*, but who seemingly do not contain a PTS1 motif, targeted to peroxisome? We hypothesized several options: (i) That they have a non-canonical PTS1 that can still bind to Pex5. (ii) That they do not contain a PTS1, but can “piggyback” on a partner protein that contains a PTS1, as has been previously shown for Mdh2 (Gabay-Maskit et al., 2020). (iii) That they bind Pex5 on a different interface (PTS1-independent), as shown for a few peroxisomal matrix proteins, the most characterized of which is Pox1 (Kempiński et al., 2020; Rymer et al., 2018; van der Klei and Veenhuis, 2006).

To unravel which of the above options is relevant for each protein, we first systematically dissected which Pex5 interface they rely on, the PTS1 or Pox1-binding interfaces. To do this we created two strains each with a point mutation in the *PEX5* gene. The first, N393D, is in the PTS1-binding pocket and weakens the interaction of Pex5 with PTS1 proteins (Klein et al., 2002). The second, Y253N, weakens the interaction of Pex5 with Pox1 and possibly other proteins that interact with Pex5 on the same interaction surface (Klein et al., 2002). Using known cargos we validated that these mutations allow us to microscopically differentiate between the two binding options (Fig. S3C and S3D). We introduced the two mutant forms of *PEX5*into the N’ GFP tagged peroxi-ome strain collection (Fig. 3A). Surprisingly we found that the targeting of all 22 novel matrix proteins depends on the PTS1 binding site of Pex5 (Data S1 and an example in Fig. 3B). Thus far, we hypothesized that the identified matrix proteins either have a new type of PTS1 or that they piggyback on a PTS1 protein.

To explore potential new PTS1 motifs, we used molecular dynamics (MD) simulations of Pex5/peptide complexes. In short, we used the available experimental structures of human Pex5/cargo complexes (Stanley et al., 2006; Fodor et al., 2012) to construct complexes of the previously modeled yeast Pex5 PTS1-binding domain (Gabay-Maskit et al., 2020), with short peptides of six C’ amino acids from known and new, putative, cargo proteins. The constancy of the peptide backbone H-bonds throughout 180nanosecond MD trajectories was used to estimate binding stability (Data S2). For the 22 known cargos of Pex5 that have a canonical PTS1, we found that the most stable H-bonds are those formed by the backbone atoms of peptide residue −1 (most C’), and residue −3. We used these to provide a measure for identifying likely peptide cargos (Fig. S4A box plot and Table S2). We applied the ranges of average stabilities for positions −1 and −3 of the known Pex5 cargos to test the MD trajectories of 12 of the newly-identified proteins, six of them previously predicted by sequence to be Pex5 cargos (Notzel et al., 2016), and six that were not previously predicted but had similarities to PTS1 sequences (Table S3). Our results predict that some of them can potentially bind to Pex5, interacting with the PTS1 binding pocket at a level similar to that of the 22 known PTS1 cargos.

To experimentally verify the binding predictions, we fused several of the newly predicted motifs to GFP, genomically integrated them into the yeast genome at a locus not affecting cell growth, and tested whether they are sufficient to support peroxisomal targeting. While most motifs were not sufficient to mediate peroxisomal localization when fused to GFP (Fig. S4B), the last 10 amino acids of Yhl045w, an uncharacterized protein, were sufficient to target GFP to peroxisomes, in a Pex5 and PTS1-binding site-dependent manner (Fig. 3C). Hence, we decided to name Yhl045w Pxp3 (<u>P</u>ero<u>x</u>isomal <u>p</u>rotein 3). Pxp3 motif was not only sufficient but also necessary for peroxisomal targeting since a C’ fusion of a short HA tag prevented it from acting as a targeting signal (Fig. S4C). Indeed, a representative frame from the MD simulations for the Pex5/Pxp3 peptide complex shows that the Pxp3 PTS1 can nicely fit into the PTS1-binding pocket (Fig. S4D).

Experimental structures of Pex5 in complex with PTS1 cargos show that the positively charged PTS1 residue in position −2 is located in a shallow, mildly negatively charged depression within the binding cavity, but its charged end makes only water-mediated H-bonds with Pex5 (Fodor et al., 2012; Stanley et al., 2006). A similar shallow, mildly negative depression is seen in modeled yeast Pex5 where the tyrosine (Y) side chain in position −2 of Pxp3 can be accommodated, making numerous contacts including water-mediated contacts that involve the OH group. To experimentally demonstrate direct binding between the potential new PTS1 and Pex5 protein and to calculate the binding affinity we used fluorescence anisotropy, which can infer precise dissociation constants (Kd) for the protein-peptide interactions (Rosenthal et al., 2020). Indeed, this assay demonstrates that purified yeast Pex5 can directly interact with the Pxp3 targeting peptide as strongly as it binds the PTS1 of a known cargo protein, Idp3 (Fig. 3D). Overall, our results confirm that Pxp3 has a *bone fide* unique PTS1, with a Y residue in position −2. This extends the consensus sequence for PTS1 motifs in yeast (Fig. 3E). Interestingly, in plants, a tyrosine residue in position −2 was also shown to allow targeting of a reporter protein to peroxisomes (Lingner et al., 2011), supporting this finding.

Having identified a new PTS1 protein, we sought to test whether the rest of the matrix proteins, that rely on the Pex5 PTS1-binding domain but do not seem to contain a PTS1, are piggybacking on a known PTS1 protein. We designed a microscopic screen in which we recorded the peroxisomal localization of each N’ GFP tagged matrix protein on the background of deletion of each of the known PTS1 proteins. While this analysis worked well in identifying the positive controls of known piggybacking cases, we were surprised to find that no deletion of a PTS1 protein affected the localization of any of the newly-identified peroxisomal matrix proteins (data not shown). This, alongside their lack of PTS1, opens an unexpected possibility that these proteins directly rely on the PTS1-binding interface of Pex5 for their peroxisomal localization without having a *bona fide* PTS1 motif.

To test for a possible direct interaction with Pex5, we chose to focus on the glutathione reductase Glr1 as a case study. Glr1 was previously shown to be a cytosolic protein with an alternative start site that generates a Mitochondrial Targeting Sequence (MTS), which targets a fraction of the protein to mitochondria (Outten and Culotta, 2004). Our whole-proteome screened showed that when the cytosolic isoform of Glr1 is N’ fused to GFP, it is partially targeted to peroxisomes in a PTS1-dependent manner (Fig. S5A).

To make sure that the peroxisomal localization is not simply resulting from interference of the GFP tag, we tested whether untagged Glr1 is indeed active in peroxisomes. Following subcellular fractionations of cells using a density gradient, we detected the peroxisomal- and mitochondrial-enriched fractions, by the activity of Fox2 (<u>F</u>atty acid <u>Ox</u>idation 2) for peroxisomes and fumarase (Fum1) for mitochondria. Measuring the activity of glutathione reduction clearly demonstrates that native Glr1 is active in both peroxisomes and mitochondria (Fig. 3F).

While all the above predictions clearly showed that Glr1 does not contain a PTS1-like sequence we verified this experimentally in two ways (i) we demonstrated that it can be targeted to peroxisomes when its most C’ is obstructed by a tag (Glr1-mNeonGreen) which is not possible for PTS1 proteins (Fig. 3G) and (ii) we measured that, unlike other PTS1 motifs, the last 10 amino acids of Glr1 are not sufficient to target GFP to peroxisomes (Fig. S5B). Hence, Glr1 was a good candidate to measure PTS1 independent binding to Pex5.

We, therefore, expressed Pex5 and Glr1 (either individually or together) in *Escherichia coli* and tested whether they can be pulled down together *in vitro*. Our assay showed that when expressed together, Glr1 was pulled down with Pex5, which indicates that the two proteins directly interact (Fig. 3H and S5C). Taken together, we speculate that Glr1 represents a whole set of proteins that have a non-canonical targeting dependency on the PTS1-binding domain of Pex5, despite lacking a PTS1 motif.

Our results open a new perspective on the complexity of peroxisomal targeting and encourage a more focused endeavor to discover the mechanistic properties underlying it.

### Systematic metabolomic analyses of peroxi-ome mutants provide clues for new enzymatic functions in peroxisomes

One of our goals in uncovering the full peroxi-ome was to extend our knowledge of the variety of peroxisomal metabolic activities. To realize this we took an unbiased approach to uncover more metabolic roles of peroxisomes and performed a large-scale metabolomic analysis for the entire peroxi-ome. For each gene of a peroxisomal protein, we analyzed the effect of its deletion or over-expression on the yeast metabolome (thousands of chemical compounds found in the yeast cell). This experiment was performed in two growth conditions – in media containing glucose or in media containing oleate (Fig. 4A and Tables S4 and S5). This resulted in over 400 profiles of mutants and a rich source of information about the effect of these mutants on cellular metabolism.

**Figure 4.**
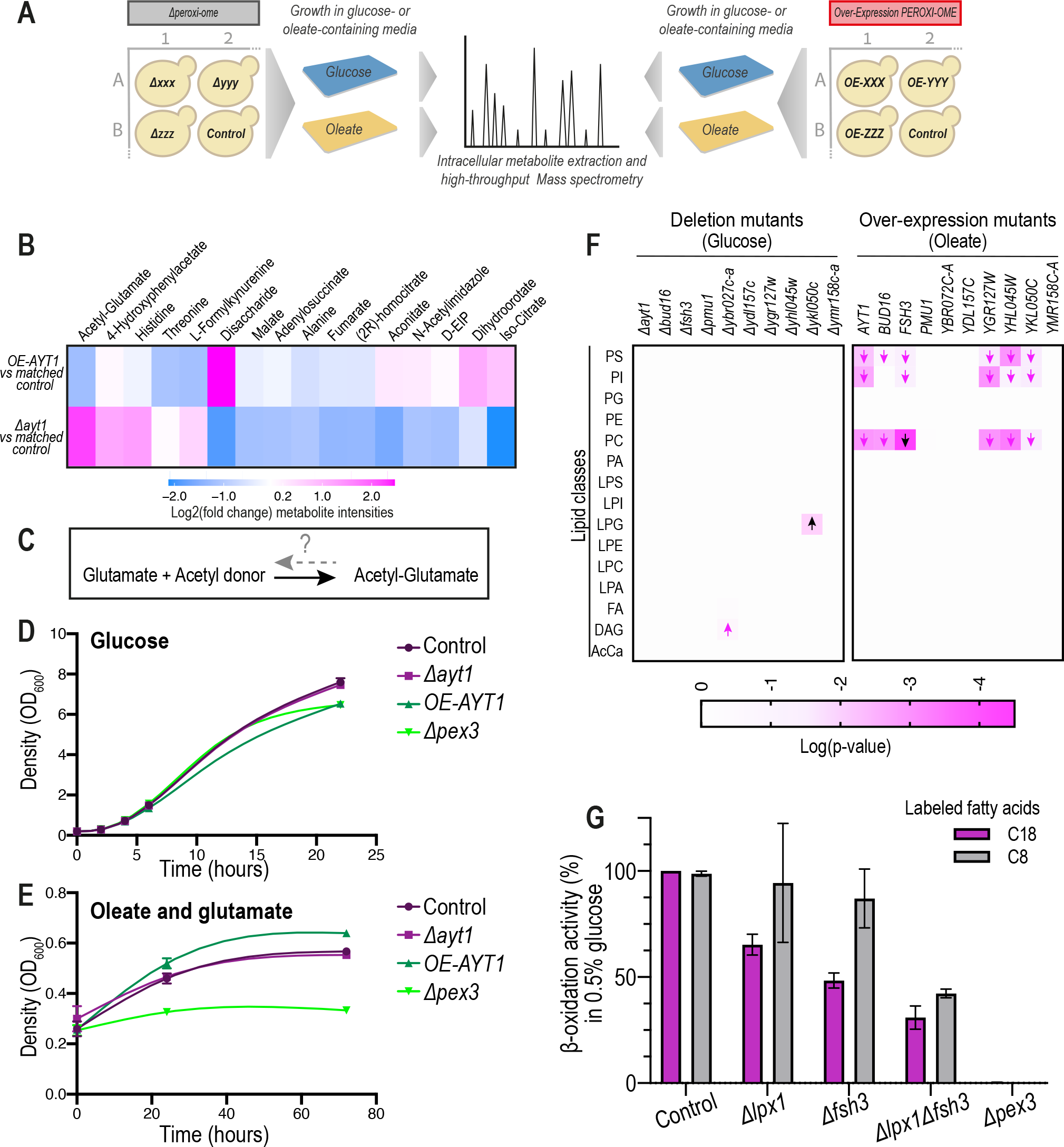
Systematic metabolomic analyses of peroxi-ome mutants provide clues for new enzymatic functions in peroxisomes. (A) Large-scale metabolomic analysis of peroxi-ome mutants (either over-expression or deletion of each gene) was performed in both glucose and oleate-containing medium to uncover additional peroxisomal metabolic functions in an unbiased manner. Raw data is in Tables S4 and S5, circular dendrograms of all conditions are in Fig. S6. (B) Metabolomic analysis focused on strains with over-expression or deletion of the *AYT1* gene shows a significant reduction or accumulation, respectively, in acetyl-glutamate compared to the matched control strain of each mutant. Only metabolites changing significantly in at least one of the two conditions are presented. D-EIP is D-erythro-1-(Imidazole-4-yl) glycerol 3-phosphate (C) Acetyl-glutamate can be generated either spontaneously in high concentrations of acetyl-CoA and glutamate, or by Arg2 or Arg7. However, it is not known how it is catabolized. (D) A growth assay in a condition that does not induce high levels of acetyl-CoA in peroxisomes (glucose) shows that a strain over-expressing *AYT1* grows to a lower density compared to the control strain. (E) A growth assay in a condition that does induce high levels of acetyl-CoA in peroxisomes (oleate as a sole carbon source) and relies on glutamate as a nitrogen source demonstrates that in conditions expected to give rise to high levels of acetyl-glutamate, the *AYT1* overexpressing strain grows faster and to a higher density than the control. (F) Lipidomic analysis on mutants of ten newly-identified peroxisomal proteins whose molecular function in the yeast cell is putative or unknown shows that cells over-expressing *FSH3* had the most significant change in all conditions compared to the control strain, with a reduction of PC (raw data is in Table S7). *Δykl050c* shows the most significant change in glucose conditions, with an increase of LPG lipids. Additional conditions are represented in Fig. S7A. Arrows indicate the directionality of the fold-change. Black arrows are the most significant changes in each condition. PS, phosphatidylserine; PI, phosphatidylinositol; PG, phosphatidylglycerol; PE, phosphatidylethanolamine; PC, phosphatidylcholine; PA, phosphatidic acid; ‘L’ indicates a ‘lyso’ phospholipid; FA, fatty acid; DAG, diacylglycerol; AcCa, acyl-carnitine (G). A β-oxidation activity assay of *Δlpx1, Δfsh3*, and *Δlpx1Δfsh3* strains supplemented with labeled 8 carbon- or 18 carbon-free fatty acids in media supplemented with 0.5% glucose shows a significant reduction in β-oxidation activity compared to the wild type and the two single mutants, suggesting an overlapping role for Fsh3 with Lpx1.

We used hierarchical clustering to uncover the relationship between the metabolomic fingerprints of the different mutants grown on each condition, and applied Gene Ontology (GO) enrichment analysis for the biological processes of known peroxisomal genes in each cluster (Fig. S6A-D). This functional clustering for mutants with similar metabolome profiles provides clues as to the functions of both known and newly identified peroxisomal proteins.

When zooming into specific metabolites, we identified the previously reported activity of several newly-identified peroxisomal proteins, such as a reduction in 3−hydroxykynurenine in the deletion of the kynurenine 3-monooxygenase, Bna4, an increase in glutathione disulfide in the absence of Glutathione reductase, Glr1, and the accumulation of pyridoxine in the absence of the putative pyridoxal kinase, Bud16 (Fig. S6E-G).

Confident that we can detect the changes in metabolites for known enzymes, we next focused on enzymes that contain protein domains predicted to perform specific metabolic activities, although their exact enzymatic activity was not yet defined. An example of such a protein is Ayt1 (Acetyltransferase 1), which was predicted to be an acetyltransferase by sequence similarity to the *Fusarium sporotrichioides* acetyltransferase, Tri101, (Alexander et al., 2002), but whose molecular function in baker’s yeast was never studied. Further metabolomics focused on the deletion and over-expression of *AYT1* revealed a significant change in the levels of acetyl-glutamate compared to the equivalent control strain (Fig. 4B and Table S6). This observation is interesting because glutamate is an important metabolite in the redox shuttle that regenerates NAD^+^ for the course of β-oxidation in peroxisomes (Rottensteiner and Theodoulou, 2006) and because the acetyltransferase domain could recognize acetyl-glutamate.

In yeast, acetyl-glutamate is generated by two mitochondrial enzymes, Arg2 and Arg7, and can also form non-enzymatically in conditions of high concentrations of acetyl-CoA and glutamate (Dercksen et al., 2016) such as those found in peroxisomes. However, how acetyl-glutamate is catabolized in yeast is not known. We hypothesized that Ayt1 may be important to regenerate glutamate from acetyl-glutamate by either using an acetyltransferase or esterase activity (Fig. 4C). To test this hypothesis in a physiological context, we performed a growth assay for strains with either a deletion or over-expression of *AYT1* in a condition that can elevate the amount of intracellular acetyl-glutamate. This media contains glutamate as the nitrogen source, in combination with oleate as the sole carbon source, that, upon β-oxidation, generates acetyl donors in peroxisomes, which can react with glutamate and increase acetyl-glutamate levels (Dercksen et al., 2016). Interestingly, while in the glucose condition, an over-expression (OE) of *AYT1* slowed yeast growth compared to the control strain (Fig. 4D), in the oleate and glutamate condition, it grew faster and to a higher density than the control (Fig. 4E). This result is in line with the metabolomic analysis, which altogether suggests that Ayt1 may catabolize acetyl-glutamate and facilitate the growth under oleate conditions.

Since our metabolomic analysis exposed potential new peroxisomal functions, we wanted to examine the involvement of several newly-identified peroxisomal proteins in lipid metabolism, as one of the hallmarks for peroxisome function is the degradation of fatty acids. We, therefore, performed lipidomic analysis for strains with either a deletion or an over-expression in one of the 10 newly-identified peroxisomal proteins whose molecular function in the yeast cell was unknown or putative according to the *Saccharomyces*Genome Database (SGD). To get a clearer view of the overall changes in the lipidomic profile of the strains, we grouped different lipids according to their classes (Table S7, Fig. 4F, and Fig. S7A). Intriguingly, the most significant change in all conditions was observed for the over-expression of Fsh3 in oleate growing cells, causing a reduction in phosphatidylcholine (PC) (Fig. 4F). Fsh3 is a putative lipase, that belongs to the family of serine hydrolases (Baxter et al., 2004). Fsh3 was predicted by sequence to contain a PTS1 motif (Notzel et al., 2016), a prediction supported by our MD computations (Figure S4A). Moreover, in a functional proteome assay in which serine lipid hydrolases were pulled down by labeled inhibitors, Fsh3 was detected in the peroxisomal fraction (Ploier et al., 2013). Indeed, we observed GFP-Fsh3 compartmentalized in the peroxisomal matrix (Fig. 2E). We confirmed that Fsh3, which contains a unique PTS1 sequence (G in position −3), is indeed a PTS1 dependent cargo by fusing its last 10 amino acids to GFP and showing that it is enough to mediate a co-localization of the GFP protein with a peroxisomal marker in a Pex5-dependent manner (Fig. S7B).

To check whether Fsh3 has an overlapping function with the only peroxisomal lipase characterized to date, Lpx1 (Lipase of Peroxisomes 1) (Thoms et al., 2008), we measured the β-oxidation activity of cells lacking either one of the two enzymes, *Δlpx1* and *Δfsh3*, or both enzymes *Δlpx1Δfsh3*. As expected, when cells grew in oleate, none of the deletions affected β-oxidation activity (Fig. S7C), plausibly due to high levels of free fatty acids in the media which would not require lipase activity. However, when cells were grown in a medium that is low in carbon (supplemented with only 0.5% glucose and thus would result in first storage of lipids as tri-acylglycerols before sugar became sparse and only then a breakdown of the phospholipids that would require a lipase activity) the double mutant *Δlpx1Δfsh3* showed a significant reduction in β-oxidation activity compared to the wild type and the two single mutants *Δlpx1* and *Δfsh3* (Fig. 4G). This effect was observed regardless of the labeled fatty acid that was added, the medium-chain fatty acid C8 or the long-chain fatty acid C18. Our findings alongside studies showing Fsh3 is a putative lipase (Baxter et al., 2004; Ploier et al., 2013), suggest that Fsh3 is a newly-identified PTS1-containing peroxisomal lipase.

A deeper inspection of previous studies on putative lipases (Ploier et al., 2013) brought to our attention that Ykl050c possesses a clear lipase activity, however, its cellular localization was never clearly determined. Our work uncovers also Ykl050c as part of the peroxi-ome and the lipidomic results show that *Δykl050c* had the most significant effect on the yeast lipidome when cells grew in glucose-containing media. This strain showed significantly higher levels of lyso-phosphatidylglycerol (LPG) (Fig. 4F). The identification of Ykl050c as a peroxisomal matrix protein (Fig. 2D) alongside our lipidomic results and it’s clear lipase activity as previously reported (Ploier et al., 2013) propose that Ykl050c is an additional newly-identified peroxisomal lipase, hence we named Ykl050c, Lpx2 (Lipase of Peroxisomes 2).

In summary, we propose that the peroxisomal matrix may contain three peroxisomal lipases (Lpx1, Lpx2, and Fsh3) that potentially act on different lipid substrates. Overall, our systematic approach provided many clues for new metabolic functions in peroxisomes and suggests an activity for several new enzymes.

### Uncovering peroxisomal targeting of GID complex subunits places peroxisomes as regulators of gluconeogenesis

The establishment of a comprehensive compendium of peroxisomal proteins in an unbiased and non-hypothesis-driven manner brought to light peroxisomal localization of several well-known proteins that would not have been suspected to be in peroxisomes. We were particularly intrigued by the co-localization of two Glucose Induced degradation Deficient (GID) complex subunits with peroxisomes when cells are grown in oleate-containing media (Table S1). The GID complex is a well-known and highly studied ubiquitin ligase that regulates glucose homeostasis, a fundamental cellular process tightly controlled by two opposing metabolic pathways - the breakdown of glucose through glycolysis and the regeneration of glucose by gluconeogenesis. These two metabolic processes share most enzymes, however, few steps are irreversible (Melkonian et al., 2020). Gluconeogenesis utilizes unique enzymes that circumvent irreversible steps of glycolysis, one of them is mediated by the enzyme fructose-1,6-bisphosphatase (Fbp1) (Fig. 5A).

**Figure 5.**
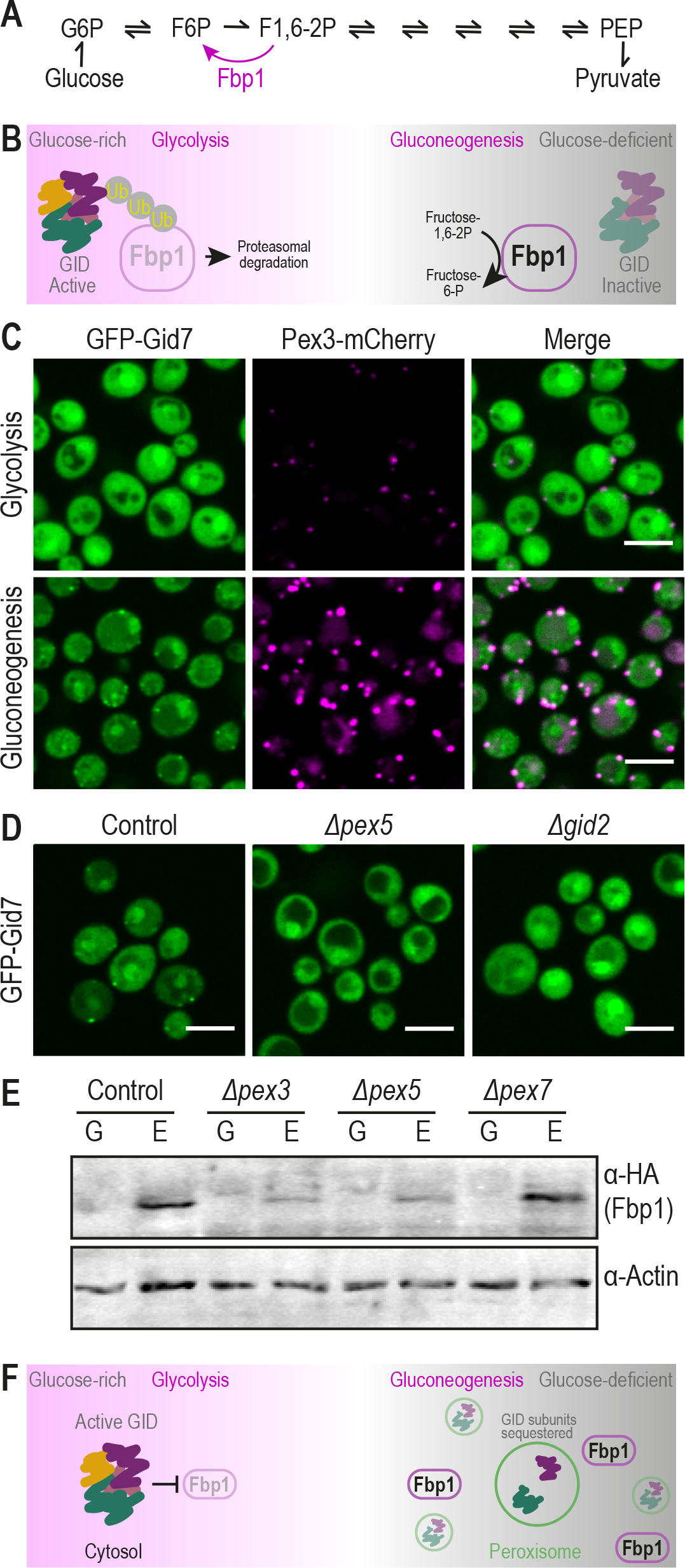
Uncovering peroxisomal targeting of GID complex subunits places peroxisomes as regulators of gluconeogenesis. (A) Gluconeogenesis utilizes unique enzymes that circumvent irreversible steps of glycolysis, one of them is the Fructose-1,6-bisphosphatase Fbp1. (B) To ensure that gluconeogenesis does not work during glucose replete conditions, the GID complex ubiquitinates gluconeogenic enzymes in glucose-enriched conditions and marks them for proteasomal degradation. In gluconeogenic conditions, the subunit that provides substrate selectivity, Gid4, is downregulated, enabling the upregulation of necessary gluconeogenic enzymes. (C) GFP-Gid7, a subunit of unknown function of the GID complex, is targeted to peroxisomes in gluconeogenic conditions (also in Fig. S9A). GFP-Gid2, which was previously shown to co-localize with peroxisomes, shows a similar phenotype (Fig. S9B). (D) The peroxisomal localization of GFP-Gid7 is dependent on both *PEX5* and *GID2*, implying that the targeting of Gid7 is mediated in the context of the complex or a sub-complex. (E) Western blot analysis of Fbp1-HA levels in mutants *Δpex3* (no peroxisomes), *Δpex5* (abolished targeting of GID subunits to peroxisomes), and *Δpex7* (no β-oxidation due to loss of Pot1 targeting) demonstrates that two hours after the transition from glucose to ethanol (gluconeogenic conditions), control and *Δpex7* cells upregulated Fbp1-HA levels, while for *Δpex3* and *Δpex5* cells Fbp1-HA levels remain low. Antibodies were used against the HA tag (for Fbp1) and Actin as a loading control. (F) Peroxisomes regulate Fbp1 levels in the transition to gluconeogenic conditions, plausibly due to partial sequestration of GID complex subunits in the peroxisomal matrix. For all micrographs, a single focal plane is shown. The scale bar is 5 µm.

The GID complex is active during glucose-rich conditions when it polyubiquitinates gluconeogenic enzymes and marks them for degradation by the proteasome (Fig 5B) (Menssen et al., 2012; Regelmann et al., 2003). Hence, it is not surprising that we detect both GFP-Gid7 and GFP-Gid2 mainly localized to the cytosol in glucose-rich conditions (Fig. 5C). Interestingly, however, GFP-Gid2 and GFP-Gid7 partially co-localized to peroxisomes when conditions become gluconeogenic, i.e. media with non-fermentable carbon sources like oleate, ethanol, and glycerol, or growth during stationary phase, when glucose is depleted (Fig. 5C, S8A, and S8B).

Moreover, Gid2 and Gid7 were not only localized to the surface of peroxisomes but were sequestered within its matrix in gluconeogenic conditions (Fig. S8C) in a manner dependent on Pex5 (Table S1). Although Gid7 and Gid2 do not interact directly in the complex (Menssen et al., 2012) we found that GFP-Gid7 was also dependent on Gid2 for proper targeting to peroxisomes (Fig. 5D) implying that the targeting of Gid7 is mediated in the context of the complex or a sub-complex. However, we could not observe other subunits inside peroxisomes using tagging and fluorescence microscopy.

What would GID subunits be doing in peroxisomes? We first assayed if GID subunits had a role inside peroxisomes but found no major effects for *Δgid2* or *Δgid7* on the levels and localization of peroxisomal matrix proteins using fluorescence microscopy (data not shown). We hence hypothesized that the possible role of peroxisomes is to sequester the subunits away from the cytosol. It was previously shown that in gluconeogenic conditions, Gid4 (also called Vid24), the subunit that provides substrate binding and selectivity, is degraded (Menssen et al., 2018), enabling the upregulation of the necessary gluconeogenic enzymes. Targeting of GID subunits to peroxisomes in gluconeogenic conditions may therefore form an additional layer of regulation on the function of this complex. To test this hypothesis, we created strains mutated in peroxisome biogenesis and checked the levels of Fbp1 in the transition from glucose to ethanol. We observed that 2 hours after the transition to ethanol, the control cells already managed to upregulate the levels of Fbp1-HA. However, in *Δpex3* cells that lack peroxisomes (Höhfeld et al., 1991) and hence have GID complex members still cytosolic, and in *Δpex5* cells in which the GID subunits are not targeted to peroxisomes, Fbp1-HA levels remained low. This effect was specific and not a by-product of reduced fatty acid catabolism since the deletion of *PEX7*, the peroxisomal targeting factor that is not mediating the targeting of GID subunits (Table S1), did not affect Fbp1 levels (Fig. 5E). This demonstrates that fully functional peroxisomes, as well as Pex5-dependent targeting, are regulating Fbp1 levels in the transition to gluconeogenesis, plausibly due to partial sequestration of GID complex subunits in the peroxisomal matrix (Fig. 5F).

## Discussion

Defining the extent of the peroxi-ome and its multifaceted functionality has been a focus and a challenge of the field for the past 70 years. Indeed, tens of proteins were identified so far in yeast peroxisomes and many of them functionally studied in great detail (Chen and Williams, 2018). Building on the assumption that more remains to be uncovered, we used a high-content screening approach and expanded the protein count of peroxisomes by ∼40%. To date, this is the most comprehensive inventory of peroxisomal proteins that can now provide a broad basis for systematic investigation of peroxisomes, allowing an in-depth understanding of how defects in peroxisomes lead to diseases. Indeed, over 60% of the yeast peroxi-ome have an established human homolog (Table S1) (Cherry et al., 2012).

Why were these proteins never assigned to peroxisomes in the past? First, previous proteomic approaches relied on the enrichment of proteins in certain fractions reducing the chances to clearly classify dual localized proteins correctly. In addition, many of the newly described peroxisomal proteins are expressed at low levels in the conditions that were previously tested. This, in combination with the fact that most of the proteins we found do not contain a canonical targeting sequence, hence could not have been detected by sequence analysis, exemplifies the sensitivity and novelty of our systematic imaging approach.

We believe that more peroxisomal proteins may be identified using imaging approaches. For example, proteins whose N’ is important for their peroxisomal targeting, must in the future be examined by fusing their C’ to a fluorophore. Moreover, some proteins are low in abundance or have only a small peroxisomal fraction. Hence, expressing their encoding genes under an even stronger promoter than *NOP1pr*, or visualizing them under different growth conditions, could expose their peroxisomal localization. Still, having this first comprehensive list enables us to leverage our extensive peroxi-ome view to perform systematic analyses never before possible on an organelle-wide level. These, alongside dedicated follow-up experiments, have already enabled the discovery of several, fundamental, concepts in peroxisome biology ranging from targeting to metabolic to regulatory mechanisms.

First, we uncovered a striking phenomenon whereby all 22 newly-identified matrix proteins relied not only on Pex5 but specifically on the PTS1 binding cavity of Pex5, although most of them did not include a PTS1 motif. In addition to the fact that these proteins do not seem to piggy-back on other PTS1 proteins, it is possible that (i) They interact directly with Pex5, as we showed for Glr1, in a mechanism yet to be studied (ii) The presumed C’ sequence of the protein as presented by the yeast genome sequencing project, does not, in fact, represent the actual C’ in cells (Schwartz and Sherlock, 2016). This can be either due to sequencing or annotation errors or due to biological mechanisms. For example, in mammalian cells, it was shown that a translational readthrough can result in the creation of a PTS1 motif for a small fraction (1.8%) of the Lactate Dehydrogenase B (LDHB) protein (Schueren et al., 2014) and a small fraction (4%) of Malate Dehydrogenase 1 (MDH1) protein (Hofhuis et al., 2016). Other mechanisms to expose a cryptic PTS1 motif include an alternative splicing (Freitag et al., 2012) and translational frameshift (Malagnac et al., 2013). Hypothetically, any type of manipulation – from cleavage to post-translational modification can result in a modified C’, which may lead to a creation of a PTS1-like motif. Such mechanisms to create PTS1 post-transcriptionally were never shown to occur in baker’s yeast. Having now exposed tens of cases of non-PTS1 Pex5-dependent proteins enables the investigation of, what seems to be, significant non-canonical mechanisms to target proteins into peroxisomes.

Another concept that came to light by our functional analysis, is the identification of two additional potential lipases in peroxisomes, Ykl050c (Lpx2) and Fsh3. Why would peroxisomes need to house a lipase, not to mention three of them, if the current dogma for lipid metabolism in peroxisomes involves the transport of free fatty acids or CoA conjugated fatty acids (acyl-CoAs) for β-oxidation (van Roermund et al., 2021)? A fascinating recent observation from *Arabidopsis thaliana* peroxisomes showed that peroxisomes form intraluminal vesicles with roles in fatty acid catabolism and protein compartmentalization (Wright and Bartel, 2020). In yeast, electron microscopy images of *Δlpx1* peroxisomes showed abnormal morphology with intraperoxisomal vesicles (Thoms et al., 2008). In the vacuole, the lipase Atg15 is known to degrade lipid vesicles (Teter et al., 2001; Epple et al., 2001) suggesting that peroxisomal lipases may have a similar role in degrading such intraluminal vesicles as a source for free fatty acids. Indeed, our high-resolution imaging demonstrated that Fsh3 compartmentalizes in the matrix to a specific niche. Moreover, deletions of both *Δlpx1* and *Δfsh3* reduce β-oxidation activity specifically when cells are grown in low glucose when there is no excess of free fatty acids in the cell. All of the above may suggest that the peroxisomal lipases Lpx1, Lpx2, and possibly also Fsh3, act on the peroxisomal membrane or on intraperoxisomal vesicles to release fatty acids for β-oxidation and in doing so also help to maintain normal peroxisome morphology. These recent findings suggest that lipid catabolism in peroxisomes is more complex than is currently thought and urge further research on the function of peroxisomal lipases.

Finally, the identification of peroxisomal targeting of GID complex subunits in glucose-deficient growth conditions adds a mechanistic view to the important concept of peroxisomes as carbon-source regulatory organelles. It has been well appreciated that peroxisomes are important for gluconeogenesis as they regenerate acetyl-CoA during β-oxidation. Subsequently, acetyl-CoA can be fed into the glyoxylate cycle and the citric acid cycle to produce metabolites for gluconeogenesis (Masters, 1997; Jardón et al., 2008). However, our results propose a regulatory role that goes beyond the simple provision of building blocks. In the same conditions that peroxisomes are most metabolically active, and in which Pex5 is upregulated, they also ensure the parallel essential presence of gluconeogenesis, plausibly by sequestering GID complex subunits and enabling the stabilization of a central enzyme in the pathway Fbp1.

The GID complex has been highly conserved through evolution in terms of sequence, however, in terms of functionality, it evolved a change in substrates. The human GID complex still regulates metabolism (Leal-Esteban et al., 2018) however the gluconeogenic enzymes FBP1 and Phosphoenolpyruvate Carboxy-Kinase (PCK1) are not direct targets of the GID complex (Lampert et al., 2018). Despite that, its co-evolution with peroxisomes seems to have persisted. For example, the murine GID complex was found to ubiquitinate AMP-activated kinase (AMPK) and therefore negatively regulate its function (Liu et al., 2020). AMPK is a regulator of cellular energy homeostasis - once activated it inhibits the transcription of gluconeogenic enzymes in the liver (Lochhead et al., 2000). In mouse hepatocytes, a deletion of PEX5 perturbed gluconeogenesis through an unknown mechanism that did, however, involve the AMPK activation (Peeters et al., 2011). It remains to further investigate whether peroxisomes regulate GID complex function also in higher eukaryotes and whether the regulation of gluconeogenesis occurs through AMPK degradation.

Interestingly, the mammalian GID complex was shown to target the transcription factor HBP1 for degradation, and thereby regulate cellular proliferation (Lampert et al., 2018). Among our list of new peroxisomal proteins, several proteins are known to be involved in cell cycle and genome duplication: the spindle pole body component Nud1 and its interactor Ady3; cell cycle regulators Mps1 and Tyc1; cell division regulator Afr1; the histone H3-like protein required for kinetochore function Cse4; and a component of the synaptonemal complex (involved in meiotic crossing over), Gmc2. Although all proteins were expressed under a constitutive promoter, Tyc1 co-localized with peroxisomes more robustly in glucose conditions compared to oleate. Tyc1 is an inhibitor of the Anaphase-Promoting Complex/Cyclosome (APC/C), a ubiquitin ligase that promotes different cell cycle phases (Schuyler et al., 2018). Yeast cells proliferate more rapidly when grown on glucose compared to non-fermentable carbon sources like oleate. Intriguingly, even when peroxisomes are artificially induced in glucose-containing media by the expression of engineered transcription factors important for peroxisome proliferation, this results in a growth delay (Grewal et al., 2021), implying that peroxisome function and the cell cycle are co-regulated. An interesting hypothesis to test in the future is whether Tyc1 is sequestered in peroxisomes to allow the proper function of APC/C in glucose-containing media. This also raises the much more global question of the role of peroxisomes and their activity in affecting cell cycle progression in response to metabolic changes. More globally our findings on the GID complex sequestration and the hypothesis on APC/C regulation support the idea that peroxisomes can be used to rapidly sequester regulatory proteins away from the cytosol (Reglinski et al., 2015).

In conclusion, our findings highlight that current knowledge on peroxisomes is only the tip of the iceberg. The discovery of multiple peroxisomal proteins introduces a more holistic perception of the peroxi-ome and the various enzymatic and regulatory activities that it may hold. More broadly, with the new understanding of the importance of peroxisomes in cellular and organismal physiology, we provide important insights that highlight new links between peroxisome regulatory and enzymatic function to carbon-source dependent cellular behavior.

## Methods

### Yeast strains and strain construction

All strains in this study are based on the BY4741 laboratory strain (Brachmann et al., 1998), except for strains based on CB199 (see the complete list of yeast strains and primers in Table S8). The libraries used were: (i) the yeast SWAT N’-GFP library, which is a collection of 5,457 strains tagged with GFP at their N’ and expressed under a generic, constitutive, promoter (SpNOP1pr) (Weill et al., 2018). 1/3 of the library was examined previously (Yifrach et al., 2016), and the additional 2/3 of the library was examined in this study. (ii) the yeast peroxisomal mini deletion library, and (iii) the yeast over-expression (*TEF2pr*-mCherry) library. Cells were genetically manipulated using a transformation method that includes the usage of lithium-acetate, polyethylene glycol, and single-stranded DNA (Daniel Gietz and Woods, 2002). Plasmids are described in Table S9. The pYM-based pMS555 plasmid that was originally used for the N-terminal GFP tagging (Yofe et al., 2016; Weill et al., 2018) was modified to contain the last 10 aa of Pxp3 (Fig. 3C), Gid7, Nud1, Ybr072c-a (Fig. S4B) and Fsh3 (Fig. S7B) at the C′ of the GFP sequence. These constructs were genomically integrated into the HO locus in strains containing Pex3-mCherry, with or without *pex5* deletion or *PEX5* N393D point mutation. Primers for validation of correct locus insertion were designed using the Primers-4-Yeast website (Yofe and Schuldiner, 2014).

### Yeast growth media

Synthetic media used in this study contains 6.7 g/L yeast nitrogen base with ammonium sulfate (Conda Pronadisa #1545) and either 2% glucose, 2% ethanol, 3% glycerol, or 0.2% oleic acid (Sigma) +0.1% Tween 80, with complete amino acid mix (oMM composition, Hanscho et al., 2012), unless written otherwise; when Hygromycin or Geneticin antibiotics were used, media contains 0.17 g/L yeast nitrogen base without Ammonium Sulfate (Conda Pronadisa #1553) and 1 g/L of monosodium glutamic acid (Sigma-Aldrich #G1626) instead of yeast nitrogen base with ammonium sulfate. When mentioned, 500 mg/L Hygromycin B (Formedium), 500 mg/L Geneticin (G418) (Formedium), and 200mg/L Nourseothricin (WERNER BioAgents “ClonNat”) were used.

### Yeast library preparation

To create collections of haploid strains containing GFP-tagged proteins with additional genomic modification such as a peroxisomal marker (Pex3-mCherry and Pex11-mScarlet) or different deletions *(Δpex5* and *Δpex7*) and point mutations (*PEX5 Y253N* and *PEX5 N393D*), different query strains were constructed based on an SGA compatible strain (for further information see strains table). Using the SGA method (Cohen and Schuldiner, 2011; Tong and Boone, 2006) the Pex3-mCherry query strain was crossed with 2/3 of the SWAT N’-GFP library and the other query strains were crossed into a collection of strains from the SWAT N’-GFP library containing ∼90 strains including known and newly-identified peroxisomal proteins together with controls. To perform the SGA in a high-density format we used a RoToR benchtop colony arrayer (Singer Instruments). In short: mating was performed on rich medium plates, and selection for diploid cells was performed on SD-URA plates containing query strain-specific antibiotics. Sporulation was induced by transferring cells to nitrogen starvation media plates for 7 days. Haploid cells containing the desired mutations were selected by transferring cells to SD-URA plates containing the same antibiotics as for selecting diploid cells, alongside the toxic amino acid derivatives 50 mg/L Canavanine (Sigma-Aldrich) and 50 mg/L Thialysine (Sigma-Aldrich) to select against remaining diploids, and lacking Histidine to select for spores with an A mating type. To create the peroxisomal mini deletion library, the BY4741 laboratory strain was transformed using plasmid pMS047 (Table S9) and with primers designed by the Primers-4-Yeast website for each peroxisomal gene.

### Automated high-throughput fluorescence microscopy

The collections were visualized using an automated microscopy setup as described previously (Breker et al., 2013). In short: cells were transferred from agar plates into 384-well polystyrene plates for growth in liquid media using the RoToR arrayer robot. Liquid cultures were grown in a LiCONiC incubator, overnight at 30°C in an SD-URA medium. A JANUS liquid handler (PerkinElmer) connected to the incubator was used to dilute the strains to an OD_600_ of ∼0.2 into plates containing SD medium (6.7 g/L yeast nitrogen base and 2% glucose) or S-oleate (6.7 g/L yeast nitrogen base, 0.2% oleic acid and 0.1% Tween-80) supplemented with –URA amino acids. Plates were incubated at 30°C for 4 hours in SD medium or for 20 hours in S-oleate. The cultures in the plates were then transferred by the liquid handler into glass-bottom 384-well microscope plates (Matrical Bioscience) coated with Concanavalin A (Sigma-Aldrich). After 20 minutes, wells were washed twice with SD-Riboflavin complete medium (for screens in glucose) or with double-distilled water (for screens in oleate) to remove non-adherent cells and to obtain a cell monolayer. The plates were then transferred to the ScanR automated inverted fluorescence microscope system (Olympus) using a robotic swap arm (Hamilton). Images of cells in the 384-well plates were recorded in the same liquid as the washing step at 24°C using a 60× air lens (NA 0.9) and with an ORCA-ER charge-coupled device camera (Hamamatsu). Images were acquired in two channels: GFP (excitation filter 490/20 nm, emission filter 535/50 nm) and mCherry (excitation filter 572/35 nm, emission filter 632/60 nm).

### Manual microscopy

Manual microscopy imaging was performed with the following strains: GFP-Nud1 with Pex3-mCherry or Spc42-mCherry (Fig. 1C); GFP-Last 10 aa of Pxp3 (Fig. 3C), Gid7, Nud1, Ybr072c-a (Fig. S4B) and Fsh3 (Fig. S7B); GFP-Pxp3 control GFP-Pxp3-HA (Fig. S4C); Glr1-mNeonGreen (Fig. 3G); GFP-Gid7 in different growth conditions (Fig. 5C and S8A and S8B) and with genetic manipulations (Fig. 5D). Yeast strains were grown as described above for the high-throughput microscopy with changes in the selection required for each strain (See yeast strain information in Table S8). Imaging was performed using the VisiScope Confocal Cell Explorer system, composed of a Zeiss Yokogawa spinning disk scanning unit (CSU-W1) coupled with an inverted Olympus microscope (IX83; x60 oil objective; Excitation wavelength of 488nm for GFP). Images were taken by a connected PCO-Edge sCMOS camera controlled by VisView software.

### High-resolution imaging

The collection of yeast strains with N’-GFP tagged peroxi-ome proteins and Pex11-mScarlet were transferred manually from agar plates into 384-well polystyrene plates (Greiner) for growth in SD-URA liquid media. Liquid cultures were grown in a shaking incubator (Liconic), overnight at 30°C. Then, strains were diluted to an OD_600_ of ∼0.2 into plates with S-oleate media. Strains were incubated for 20 h at 30°C to induce enlargement of peroxisomes and transferred manually into glass-bottom 384-well microscope plates (Matrical Bioscience) coated with Concanavalin A (Sigma-Aldrich). After 20 min, cells were washed three times with double-distilled water to remove non-adherent cells and to obtain a cell monolayer. The plate was then imaged in an automated inverted fluorescence microscope system (Olympus) harboring a spinning disk high-resolution module (Yokogawa CSU-W1 SoRa confocal scanner with double microlenses and 50 μm pinholes). Images of cells in the 384-well plates were recorded in the same liquid as the washing step at 30°C using a 60X oil lens (NA 1.42) and with a Hamamatsu ORCA-Flash 4.0 camera. Fluorophores were excited by a laser and images were recorded in two channels: GFP (excitation wavelength 488 nm, emission filter 525/50 nm) and mScarlet (excitation wavelength 561 nm, emission filter 617/73 nm). All images were taken in a Z-stack and using cellSens software. The best focal plane showing the “ring-like” structure of the peroxisomal membrane was chosen for defining sub-organellar localization. For presentation, images were deconvoluted using cellSens software.

### Molecular dynamics (MD) simulations

Exploring the association of a peptide with the Pex5 receptor via MD simulations would require excessively long computations. We, therefore, chose to explore the stability of Pex5-peptide complexes. Each simulation started from a Pex5-peptide complex modeled based on the experimental structure of human Pex5-cargo complexes, and it was assumed that improbable or unstable complexes would dissociate or weaken as the simulation proceeds. Two 100ns trajectories were calculated for each complex. The last 90ns of the two trajectories were analyzed together, providing estimates of selected contacts stability for a combined 180ns simulation. To estimate the effect of time on the Pex5-peptide contacts, the analyses were repeated for the last 50ns of the two trajectories, 100ns altogether.

Our analyses focused on the direct hydrogen (H)-bond interactions between the backbone of the peptide and all polar/charged side chains lining the PTS1 binding cavity of Pex5, defining an H-bond as a distance of 0.35nm or less between H-bond donors and acceptors. The analysis distinguishes between backbone oxygen atoms, which can each accept two or more H-bonds, and backbone nitrogen atoms that can each donate one H-bond. The C’ Ot atoms can accept four H-bonds and these are individually analyzed, and so are the two H-bonds that O-2 accepts. Notably, distance analyses showed that the Nε and Nη atoms of Pex5 Arg526 are often at an H-bond distance from peptide atom O-2. However, the direction of Arg526 N-H bonds is inadequate for H-bond formation therefore these contacts were not considered as H-bonds in the analyses. O-3 and O-5 form only one H-bond contact in the experimental structures but in the MD trajectories H-bonds were also formed to the sidechain of Tyr468. This residue replaces N462 in human Pex5 and being larger, it protrudes into the PTS1 binding cavity. The details of individual contacts are given in the supplementary Table S2 but the analyses considered the sum of the two contacts for each backbone oxygen. For backbone N-3 and N-5 atoms, the analysis considers only the shortest H-bond distance in each MD frame.

MD simulations for human Pex5 in complex with peptide YQSKL (entry 1FCH in the Protein Data Bank) were used as a test case. Most of the H-bond contacts seen in the experimental structure are well preserved in the MD trajectories, as detailed in Table S2. The Ot atoms make stable H-bonds with Asn378/Nδ2 (85.5% of the trajectory) and Asn489/Nδ2 (62.0% of the trajectory), as seen in the experimental structure. They also form very frequent H-bonds with the sidechains of Lys490 and Arg520, 43.6 and 93.6% of the trajectory, respectively, which in the experimental structures make only water-mediated H-bonds with the cargo. Most H-bonds of peptide residues −2 and −3 are also well preserved in the human Pex5 complex simulation (>50% of the time); less preserved is the contact N-2 with Asn524/Oδ1, maintained only 36.1% of the time. In contrast, the H-bonds formed by the backbone of residue −5 are not stable in the simulations. This result is not surprising as in the experimental structure residue Y-5 is also stabilized through contacts with a neighboring molecule related by crystal symmetry, while the MD simulation is for a single molecular complex.

Hydrogen bond stabilities throughout the MD trajectories for 22 known yeast Pex5 cargos were used to establish a plausible measure for identifying stable binders among the new peroxisomal proteins. The most stable hydrogen bonds were formed by the backbone atoms of peptide residues −1 and −3. The ranges of the averaged hydrogen-bond stabilities for each of these residues distinguished well between the known cargos and Mdh2, a known non-cargo, and thus were used to identify stable binders (Table S3). The time dependence of the H-bonds stability is minor, indicating that rapid structural changes occur in the first 10ns of the trajectory, which are not included in the analysis.

MD simulations were executed with the Gromacs package (Van Der Spoel et al., 2005). Trajectory analyses were performed with Gromacs and with programs written by M.E. UCSF-chimera (Pettersen et al., 2004) was used to model starting structures of the Pex5/peptide complexes and to produce Figure S4D.

### Pex5 protein purification for fluorescence anisotropy

Full-length Saccharomyces cerevisiae Pex5p was cloned in a petM30 vector. Pex5p was expressed in autoinduction medium (Studier, 2005), with 5 hours at 37°C and 26 hours at 20°C. Cells were harvested, resuspended in lysis buffer (50 mM Hepes pH 7.5, 150 mM NaCl, 20 mM imidazole, protease inhibitor (Roche), DNAse (Sigma), and lysozyme (Sigma)), homogenized 1 hour at 4°C and lysed by sonication. The lysate was then cleared by centrifugation and the supernatant loaded onto Ni-NTA resin. Bound proteins were washed with 50 mM Hepes pH 7.5, 750 mM NaCl, 20 mM Imidazole, and the protein was eluted with 50 mM Hepes pH 7.5, 150 mM NaCl, 250 mM Imidazole. The eluate was then dialyzed against Hepes pH 7.5, 150 mM NaCl, 0.5 mM TCEP and simultaneously digested with 1 mg of TEV-protease. Undigested protein and TEV protease were removed by a second Ni-NTA step and flow through containing Pex5p were concentrated for gel filtration (Hiload 16/60 Superdex 200 pg, GE healthcare). Relevant fractions were pooled together and the protein was concentrated, flash-frozen in liquid nitrogen, and stored at −80 °C.

### Fluorescence anisotropy

Fluorescein isothiocyanate (FITC) labeled peptides corresponding to the carboxyl-terminal 10 amino acids of Yhl045w (FITC-RKRVLGVAYL, Genscript) and Idp3 (FITC-YEDKKGMCKL, Genscript) were solubilized in water and used in the assay at a final concentration of 10 nM. A tyrosine was added at the N terminal of Idp3 for concentration determination. Measurements of fluorescence anisotropy changes were performed in black 96-well plates (Greiner) with an Infinite M1000 plate reader (TECAN) with excitation/detection at 470/530 nm. The experiment was performed in 50 mM Hepes pH 7.5, 150 mM NaCl. A concentration range from 38 µM to 120 nM (for Idp3) or 20 µM to 150 nM (for Yhl045w) was obtained by serial dilution and each concentration was measured in triplicate. Three independent experiments were performed and binding data were normalized and analyzed using Prism (GraphPad software, USA). Kinetic information was obtained by least-square fitting of a binding– saturation model with one binding site.

### Glr1 and Pex5 plasmid construction

All cloning reactions were performed by the Restriction-Free (RF) method (Unger et al., 2010). Full-length yeast *PEX5* was cloned into the expression vector pET28-bdSumo (Zahradník et al., 2019). Yeast *GLR1* was cloned into the first ORF of the expression vector pACYCDuet-1 (Novagen) including an N-terminal Flag-tag followed by a TEV cleavage site.

### Glr1 and Pex5 protein expression

*GLR1* (in *pACYCDut-FLAG-GLR1*) and *PEX5* (in *pET28-bdSUMO-PEX5*) were either individually or co-expressed in *E. coli* BL21(DE3). Expression was performed in LB medium supplemented with the appropriate antibiotics (Kanamycin and/or chloramphenicol). Expression was induced with 200μM IPTG followed by shaking at 15 °C for ∼16 hours. Cell pellets were stored at −20 °C before processing.

### Glr1 and Pex5 protein pull-down

Cells were lysed by sonication in Tris-buffered saline (TBS) buffer supplemented with 1 mM phenylmethylsulfonyl fluoride (PMSF) and 1μl/mL of protease inhibitor cocktail (Set IV, EMD Chemicals, Inc). Protein pull-down experiments were performed using Anti-His affinity Resin (#1018-25, Adar Biotech) according to the manufacturers’ recommendations. Western blot analysis was performed using THE^TM^ DYKDDDDK Tag Antibody [HRP-conjugated] (A01428, GenScript) and Monoclonal Anti-polyHistidine−Peroxidase (A7058, Sigma). Proteins were analyzed on 4-20% SurePAGE precast gels (M00657, GeneScript).

### Metabolite extraction for high-throughput metabolomics

A target OD_600_ of 1.5 was set for all metabolite extractions for peroxisomal mutant collection experiments with at least two cell doublings occurring between culture inoculation and extraction. For analysis of *AYT1* shown in figure 4, OD_600_ values between 0 and 1.5 were collected for each of the treatment conditions. Cultivations were performed in a 96-well format with starting volumes of 1.2 mL per well. OD_600_ values were measured before harvesting cells for extraction. For glucose-grown samples, cells were grown for 4 hours in synthetic defined media as described in the ‘yeast growth media’ section, including 2% (w/v) glucose. For oleate-grown samples, cells were first grown in synthetic defined media with a low concentration of glucose (0.1% w/v) for 16 hours. Then, cells were transferred to oleate-containing media (0.2% w/v) and were allowed to grow for additional two hours before harvesting. Cells were harvested by centrifugation for 1 minute at 2254 rcf. After discarding the supernatant, 150 µL of cold extraction solution (40% (v/v) HPLC-grade acetonitrile (Sigma-Aldrich: 34998), 40% (v/v) HPLC grade methanol (Sigma-Aldrich: 34885), 20% (v/v) HPLC-grade water (Sigma-Aldrich: 1153331000)) was added to each cell pellet. Extraction was allowed to proceed at −20°C for a duration of one hour in a covered container. The extracts were exposed to 1 minute of centrifugation at 2254 rcf and 100 µL of the supernatant was taken and transferred into conical 96-well plates (Huber lab: 7.1058). Plates were sealed (Huber lab: 7.0745) and placed at −80°C until the time of measurement.

### Flow injection time-of-flight mass spectrometry for high-throughput metabolomics

Mass spectrometric measurements were made using an Agilent 6550 Series quadrupole time-of-flight mass spectrometer (Agilent) through an adaptation of the method described by Fuhrer et al., 2011. An Agilent 1100 Series HPLC system (Agilent) was coupled to a Gerstel MPS 3 autosampler (Gerstel) to perform the analysis. A mobile phase flow rate of 0.15 mL/min was used, with the isocratic phase composed of 60:40 (v/v) isopropyl alcohol and water at a buffered pH of 9, with 4 mM ammonium fluoride. Taurocholic acid and Hexakis (1H, 1H, 3H-tetrafluoropropoxy–phosphazine) within the mobile phase were used to perform online mass axis correction. The instrument was run in high-resolution (4 GHz) mode, and mass spectra between 50 and 1000 m/z were collected in negative mode. Raw data files were deposited in the MassIVE repository (https://massive.ucsd.edu/). Peroxisomal deletion library data was deposited with accession code MSV000086773 in the MassIVE database (massive.ucsd.edu), over-expression (*TEF2* promoter) library data was deposited under code MSV000086775, and the focused *AYT1* analysis data was deposited under code MSV000086772. Datasets can be accessed in prepublication with login names “MSV000086773_reviewer”, “MSV000086775_reviewer”, “MSV000086772_reviewer” respectively, and password “reviewerpass”.

### Analysis of metabolomics mass spectrometry data

Centroiding of the mass spectrum, merging, and ion annotation was performed as described in Fuhrer et al., 2011. Metabolites used for ion annotation were drawn from the KEGG Saccharomyces cerevisiae metabolite library. Data normalization and analysis were performed using the Pandas package (McKinney, 2019) in Python. For the analysis of peroxisomal mutant collections, temporal drifts, as well as OD_600_ effects in ion intensity were corrected for using a LOWESS and linear regression approach respectively. Outlier samples in terms of OD_600_ at the time of sampling as well as in total ion current were discarded. Z-score transformations or fold-change calculations were applied to the normalized data. For the analysis of *AYT1* specifically, the slope of ion intensity with respect to OD_600_ was calculated and average fold-changes were calculated for *AYT1* mutants relative to WT based on those slopes.

### Growth assay for AYT1 strains

Growth assays were performed with strains based on CB199 (Oeljeklaus et al., 2012; Schummer et al., 2020) strain (see Table S8). Cells were incubated in 0.3% liquid glucose media, rotating overnight at 30 °C. Then, a total of 2 O.D600 of cells were centrifuged for 3 minutes at 3,000 g, and pellets were inoculated in 10 ml of either SD (6.7 g/L yeast nitrogen base with ammonium sulfate and 2% glucose), or S(MSG)-oleate (6.7 g/L yeast nitrogen base without ammonium sulfate, with 1 g/L monosodium glutamate, 0.1% oleate, 0.1% yeast extract, and 0.05% Tween-40). Both media contained a complete amino acid mix. Before measuring the O.D_600_ in oleate, 1 ml of cells was taken, cells were washed twice with water and resuspended in 1 ml water.

### Lipid extraction for high-throughput lipidomics

The indicated yeast strains were cultivated in synthetic defined media supplemented with amino acids and G418 (deletion strains) or NAT (over-expression strains) antibiotics. For glucose-grown samples, cells were grown for 4 hours in synthetic defined media as described in the ‘yeast growth media’ section, including 2% (w/v) glucose. For oleate-grown samples, cells were first grown in synthetic defined media with a low concentration of glucose (0.1% w/v) for 16 hours. Then, cells were transferred to oleate-containing media (0.2% w/v) and were allowed to grow for additional two hours before harvesting. Yeast were harvested by centrifugation at 2254 rcf for 1 minute and the supernatant was discarded. Lipid extraction was performed as described previously (Pellegrino et al., 2014) with some modifications. To 20 µl of the sample, 1 ml of a mixture of methanol: IPA 1:1 (v/v/v) was added. The mixture was fortified with the SPLASH mix of internal standards (Avanti Lipids). After brief vortexing, the samples were continuously mixed in a Thermomixer (Eppendorf) at 25°C (950 rpm, 30 min). Protein precipitation was obtained after centrifugation for 10 min, 16000 g, 25°C. The single-phase supernatant was collected, dried under N2, and stored at –20 °C until analysis. Before Analysis, the dried lipids were re-dissolved in 100µL MeOH:Isopropanol (1:1 v/v).

### Liquid chromatography and mass spectrometry for lipidomic analysis

Liquid chromatography was done as described previously (Cajka and Fiehn, 2016) with some modifications. The lipids were separated using C18 reverse-phase chromatography. Vanquish LC pump (Thermo Scientific) was used with the following mobile phases; A) Acetonitrile:water (6:4) with 10mM ammonium acetate and 0.1% formic acid and B) Isopropanol: Acetonitrile (9:1) with 10mM ammonium acetate and 0.1% formic acid. The Acquity BEH column (Waters) with the dimensions 100mm * 2.1mm * 1.7µm (length*internal diameter*particle diameter) was used. The following gradient was used with a flow rate of 1.2 ml/minutes; 0.0-0.29 minutes (isocratic 30%B), 0.29-0.37 minutes (ramp 30-48% B), 0.370-1.64 minutes (ramp 48-82%B),1.6-1.72 minutes (ramp 82-99%), 1.72-1.79 minutes (isocratic 100%B), 1.79-1.81 minutes (ramp 100-30% B) and 1.81-2.24 minutes (isocratic 30%B). The liquid chromatography was coupled to a hybrid quadrupole-orbitrap mass spectrometer (Q-Exactive HF-X, Thermo Scientific). A full scan acquisition in negative and positive ESI was used. A full scan was used scanning between 200-2000 m/z at a resolution of 60000 and AGC target 1e6. The maximum injection time was 100 ms.

### Analysis of lipidomics mass spectrometry data

Lipid identification was achieved using the following criteria; 1) high accuracy and resolution with accuracy within m/z within 5 ppm shift from the predicted mass, 2) Isotopic pattern fitting to expected isotopic distribution, and 3) retention order compared to an in-house database. Quantification was done using single-point calibration against the SPLASH internal standard (Avanti Lipids). Mass spectrometric data analysis was performed in Compound Discoverer 3.1 (Thermo Scientific) for peak picking, integration, and annotation. Class enrichment analysis was performed through the application of a hypergeometric test for enrichment of significantly changing lipids compared to the representation of those classes within the set of annotated lipids. Raw mass spectra were deposited with accession code MSV000086777 in the MassIVE database (massive.ucsd.edu). Access is possible for reviewers with login “MSV000086777_reviewer” and password “reviewerpass”.

### β-oxidation activity in peroxisomes

β-Oxidation assays in intact cells were performed as described previously (van Roermund et al., 1998) with slight modifications. Cells were grown overnight in media containing 0.5% glucose. The β-oxidation capacity was measured in 50 mM MES, pH 6.0 supplemented with 10 μM 1-^14^C-octanoate (C8:0) or 10 μM 1-^14^C oleate (C18:1). Subsequently, [^14^C]CO_2_ was trapped with 2 M NaOH and acid-soluble counts (ASP) were used to quantify the rate of fatty acid oxidation. Results are presented as percentages relative to the rate of oxidation of wild-type cells.

### Subcellular fractionation

Subcellular fractionation was performed as described by (Van der Leij et al., 1992). Briefly, a protoplast-free and a nuclei-free organellar pellet was loaded onto a 15-35% continuous Nycodenz gradient (111 ml), with a cushion of 50% Nycodenz (1 ml), dissolved in buffer A (5 mM MES pH 6.0, 1 mM EDTA, 1 mM KC1 and 8.5% (w/v) sucrose). After centrifugation for 2.5 hours in a vertical rotor (MSE 8×35) at 29,000*g at 4°C, the gradient was unloaded from the bottom, yielding 12 fractions.

### Enzymatic activity of Fum1, Pox2, and Glr1 in subcellular fractions

The 3-hydroxyacyl-CoA dehydrogenase activity of Pox2 was measured as described by Wanders et al., 1990. Fumarase was measured as described by van Roermund et al., 1999. Glutathione reductase activity was measured at 37°C by monitoring the absorption at 340 nm over a period of 5 minutes using a reaction medium with the following components: 1 mM EDTA, 50 mM TRIS buffer pH 7.2, 0.1% (v/v) Triton X-100, 0.1 mM NADPH, 5 mM GSSG and the sample to be analyzed. Since the reaction in the absence of enzyme does take place at an appreciable rate, the change in absorbance of a solution containing no sample was also recorded and subtracted.

### Western blot for Fbp1-HA strains

Cells were grown in glucose-containing media with appropriate selections overnight at 30° C. Then, cells were diluted to 0.2 OD_600_ in YPD (2% Peptone + 1% Yeast Extract (Formedium) and 2% glucose) and grown for additional 3 hours. At mid-log (0.5-0.8 OD_600_), a total of 3 OD_600_ cells were taken, washed with double-distilled water, and the pellet was snap-frozen in liquid nitrogen. In parallel, an additional 3 OD_600_ of cells were taken, washed with double-distilled water, and inoculated in an S-ethanol medium. Cells were incubated for 2 hours at 30° C and then washed with water, pelleted, and snap-frozen in liquid nitrogen. Samples were stored at −80° C until protein extraction.

For protein extraction, pellets were resuspended in urea lysis buffer (8 M Urea, 50 mM Tris pH 7.5, and 1:200 freshly added protease inhibitor cocktail (Merck)). 100 ul acid-washed glass beads were added, and cells were broken using a bead beater machine, for 10 minutes at 4° C. Then, each sample was added with 2.5% SDS and incubated at 45° C for 15 minutes. Samples were centrifuged at 3,000 g, 16° C for 10 minutes, and supernatants were transferred to new tubes for additional centrifugation at max speed, 16° C, for 5 minutes. Supernatants were collected and mixed with loading buffer (60 mM Tris pH 6.8, 10% glycerol, Orange G dye (Sigma), and 100 mM fresh DTT). Before loading on SDS-PAGE, samples were incubated again at 45° C for 15 minutes. Samples ran at a constant voltage of 80 and then transferred to a nitrocellulose membrane using a semi-dry Bio-Rad transfer machine. Antibodies used were anti-HA (BioLegend, monoclonal antibody raised in mouse, 1:1000 dilution) for Fbp1-HA, and Actin (Abcam, polyclonal antibody raised in rabbit, 1:1000 dilution).

### Western blot on *NOP1pr*-GFP newly-identified peroxisomal proteins

Cells were grown in glucose-containing media as described in the main text. Whole-cell protein extraction was performed by either the Urea protein extraction method (described in Fbp1-HA western blot method) or the NaOH protein extraction method. In short, for extracting proteins using the NaOH method, 3 OD_600_ of cells were incubated in 0.2 M of NaOH for 5 min, at room temperature. Following centrifugation, pellets were re-suspended with SDS-sample buffer (60 mM Tris pH 6.8, 2% SDS, 10% glycerol, 200 mM DTT, and 0.2 mM freshly added PMSF), and were boiled at 95° C for 5 minutes. Before loading on an acrylamide gel, samples were centrifuged at max speed for 3 minutes. Samples ran at a constant voltage of 80 and then transferred to a nitrocellulose membrane using a semi-dry Bio-Rad transfer machine. The antibody used was anti-GFP (Abcam, polyclonal antibody raised in rabbit, 1:2000 dilution).

## Acknowledgments

This work was supported by the ERC CoG OnTarget (864068). The robotic system of the Schuldiner lab was purchased through the kind support of the Blythe Brenden-Mann Foundation. MS is an Incumbent of Dr. Gilbert Omenn and Martha Darling Professorial Chair in Molecular Genetics. EY is supported by the Ariane de Rothschild Women Doctoral Program. We thank Renate Maier and Prof. Bettina Warscheid for kindly sharing the CB199 strain that was used for the growth assays and to Dr. Michal Skruzny for sharing the mScarlet tagging plasmid. We also thank Prof. Oren Schuldiner and Dr. Noa Dahan for critically reviewing the paper.

## Author contribution

E.Y, D.H.S, J.B, A.O, C.W.T.V.R, W.V, A.T, C.B, S.G, and U.W performed the experiments; M.E performed the molecular dynamics simulations and analyses; EY and AF performed the high-throughput and high-resolution screens; Y.P, H.R.W, R.J.A.W, M.W, N.Z, M.S, and E.Z supervised the work; M.S, E.Z, and E.Y wrote the manuscript. All authors read and gave feedback on the manuscript.

---

**Figure S1.**
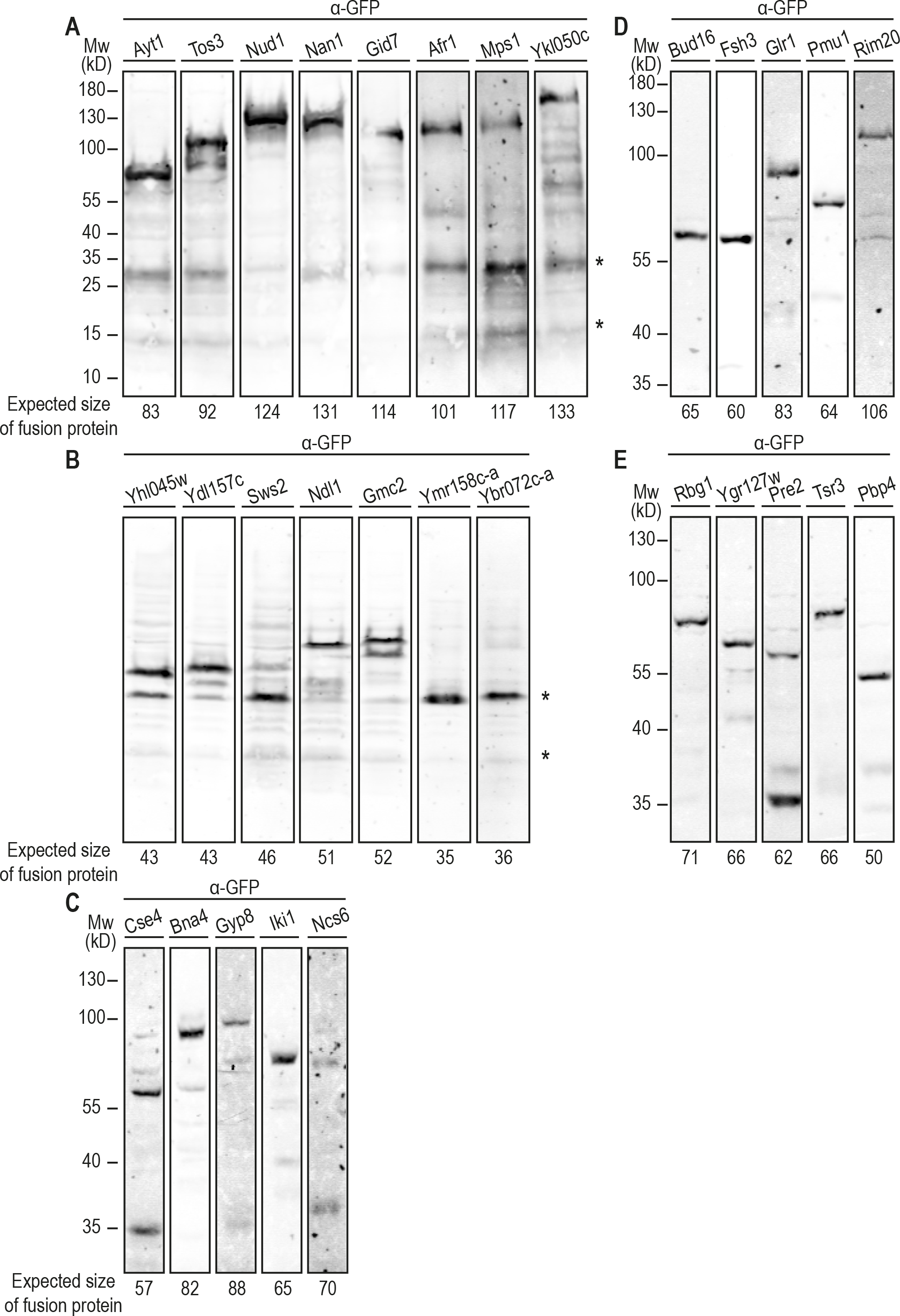
Western blot analysis of candidate peroxisomal proteins ascertain correct genomic integration. Protein samples of strains expressing each N’ GFP candidate peroxisomal protein were immunodecorated with antibodies against the GFP tag and analyzed for their expected molecular weight. Asterisks indicate non-specific bands. Tad3, Tyc1, and Ady3 were not detected. Panels (A) and (B) show analysis for proteins that were extracted by the Urea method. Panels (C)-(E) show analysis for proteins that were extracted by the NaOH method. Pre-cast 4-20% acrylamide gel (Bio-Rad) was used for panels (A) and (C)-(E). A 12.5% acrylamide gel was used for panel (B).

**Figure S2.**
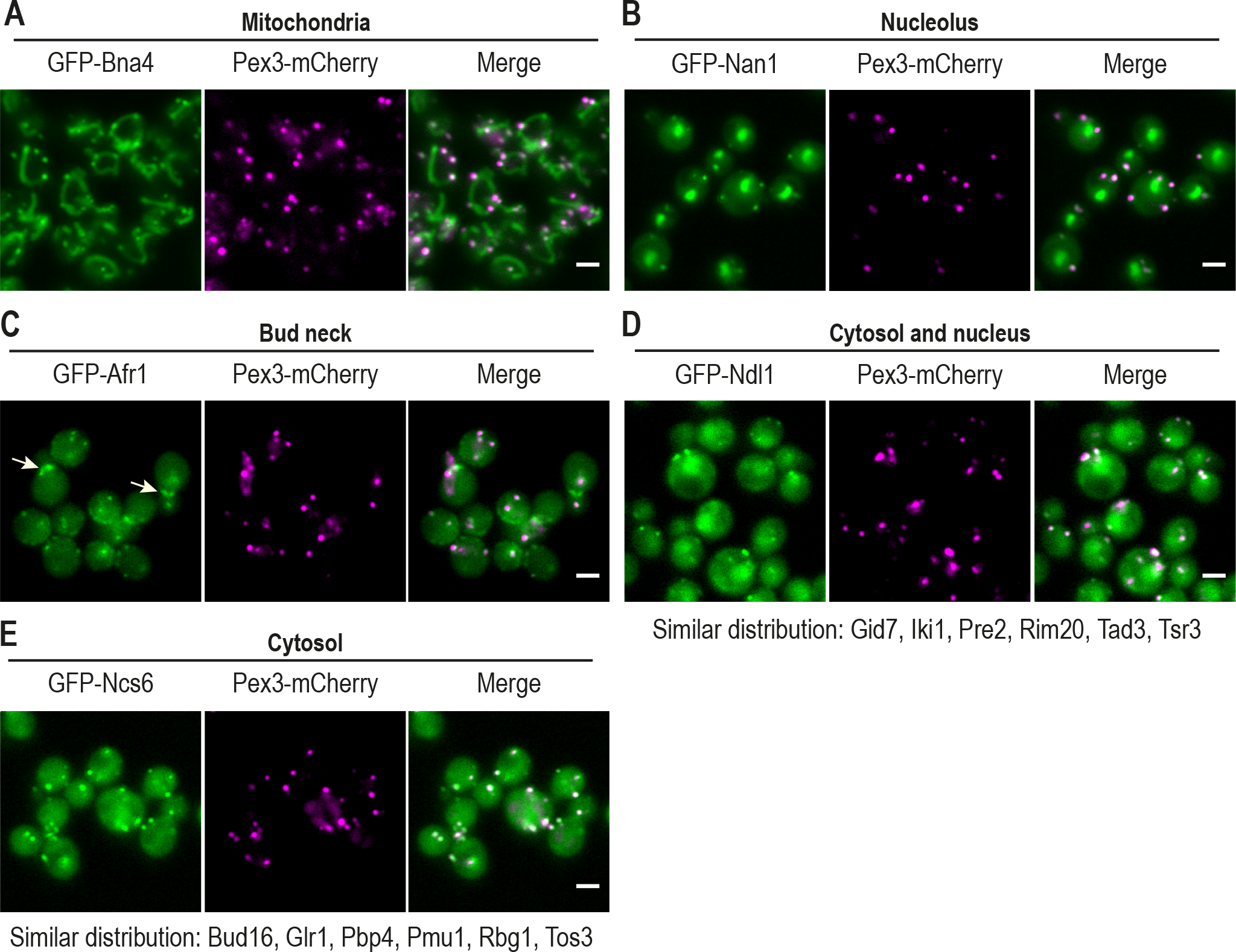
Image analysis of the newly-identified peroxisomal proteins indicates that half of them have dual cellular localization. Known localization of the newly-identified peroxisomal proteins was taken from their ‘cellular compartment’ annotation in the *Saccharomyces* Genome Database (SGD) (Cherry et al., 2012) (see also Table S1). (A) GFP-Bna4 is dually localized to mitochondria and peroxisomes. (B) GFP-Nan1 is dually localized to nucleolus and peroxisomes. (C) GFP-Afr1 is localized to the bud neck (white arrows) and peroxisomes. (D) Seven proteins are localized to the cytosol, nucleus, and peroxisomes. GFP-Ndl1 is an example from this group. (E) Seven proteins are dually localized to the cytosol and peroxisomes. GFP-Ncs6 is an example from this group. Proteins with the same distribution are listed below each image. For all micrographs, a single focal plane is shown. The scale bar is 5 um.

**Figure S3.**
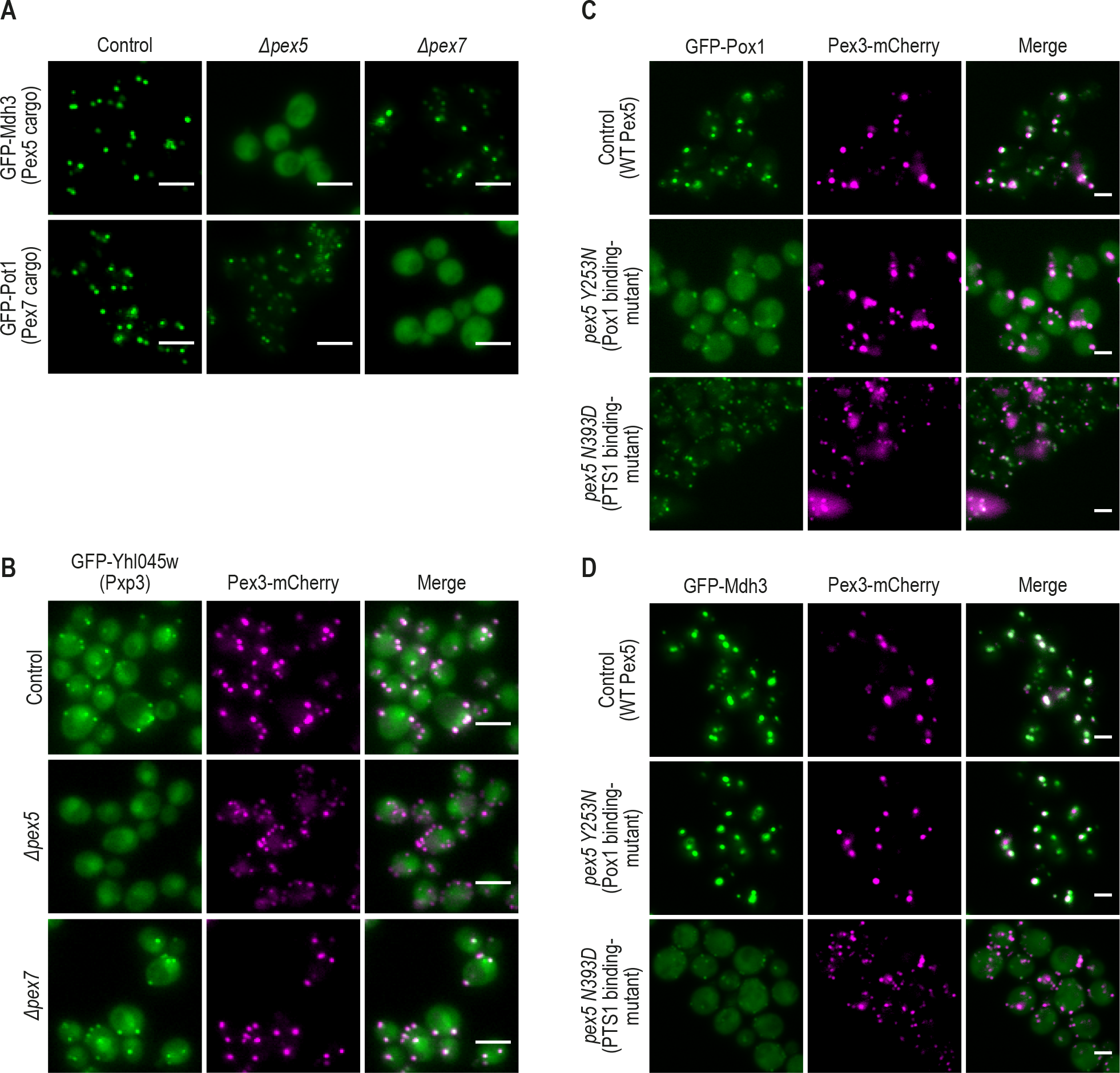
Verification of functional targeting assays using known cargo proteins. (A) Mdh3 (a Pex5 cargo protein) and Pot1 (a Pex7 cargo protein) were utilized to show expected dependencies on their specific targeting factor using fluorescence microscopy. (B) GFP-Yhl045w (Pxp3), a newly-identified peroxisomal matrix protein, represents a case for targeting dependency on Pex5, but not on Pex7. (C) GFP-Pox1 (a Pex5 cargo protein that interacts with Pex5 in a PTS1-independent manner) was used to show expected dependency upon Pex5Y253N point mutation (D) GFP-Mdh3 (a known PTS1 protein) was used to show targeting dependency on Pex5N393D (a mutation in the PTS1-binding site). For all micrographs, a single focal plane is shown. The scale bar is 5 um.

**Figure S4.**
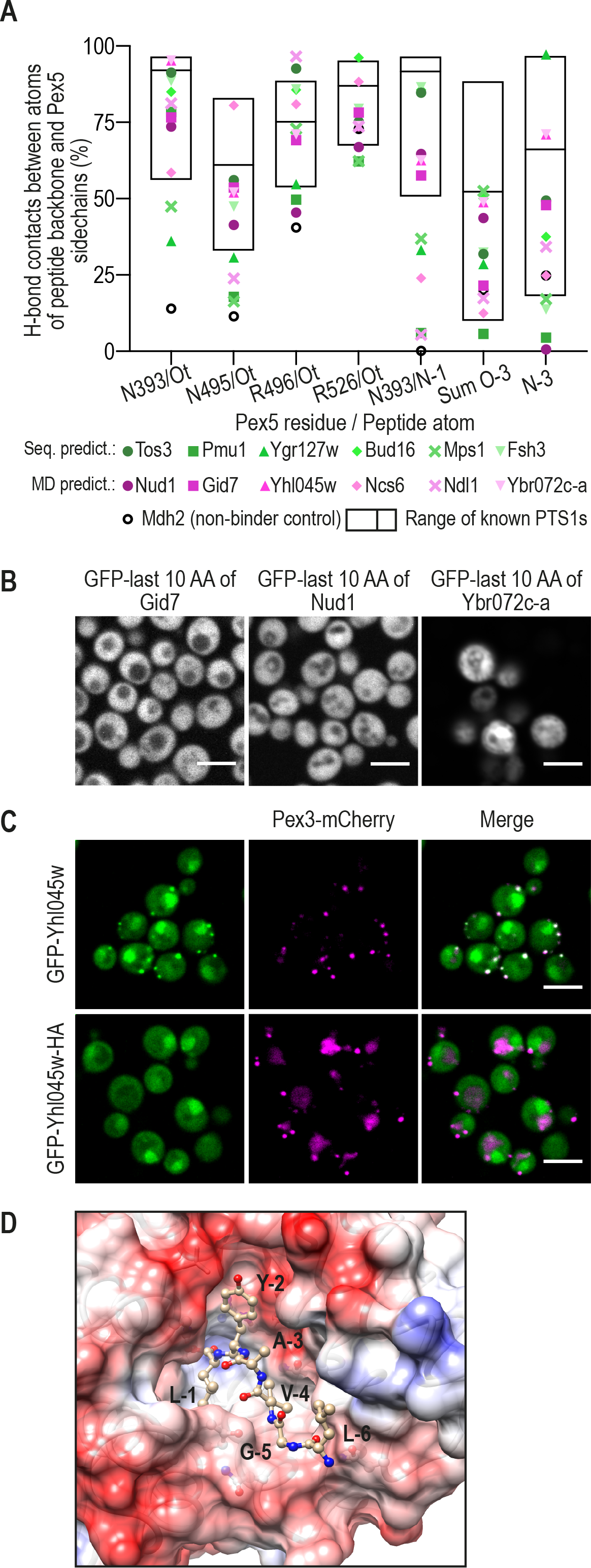
Characterization of unique PTS1 motifs. (A) Predictions for identifying potential new PTS1 motifs were made by molecular dynamic (MD) simulations. The likelihood of stable binding between Pex5 and the backbone atoms at the C’ amino acids −1 and −3 of the new, putative, cargo proteins was compared to the stability of hydrogen bonds of known PTS1 cargo proteins to Pex5 (Table S3). The ranges of hydrogen bonds stabilities for the 22 known PTS1 cargos are presented by box plots; the black lines represent averages. MD results for potential Pex5 cargos previously predicted based on their sequence are presented in green shades. Six proteins predicted by the MD analysis as cargos are presented in magenta shades. Mdh2 was used as a known Pex5 non-binder control (B) The peroxisomal targeting ability of the predicted motifs was examined by fusing the last 10 amino acids of Gid7, Nud1, and Ybr072c-a to the C’ of GFP, integrating the construct into an inert locus in the yeast genome, and imaging. Most motifs, except for the Yhl045w (Pxp3) motif (Fig. 3C), were unable to target GFP to peroxisomes (C) Blocking the C’ of GFP-Yhl045w (Pxp3) with HA tag demonstrates that Yhl045w (Pxp3) C’ is necessary for targeting to peroxisomes. (D) A representative frame from the MD simulations for the Pex5/Pxp3 complex showing the unique Pxp3 PTS1 motif nicely fit into the PTS1-binding pocket of Pex5. The side chain of tyrosine (Y) in position −2 of Pxp3 can be accommodated in a shallow, mildly negatively charged depression within the binding cavity, making numerous contacts including water-mediated contacts that involve the OH group. For all micrographs, a single focal plane is shown. The scale bar is 5 um.

**Figure S5.**
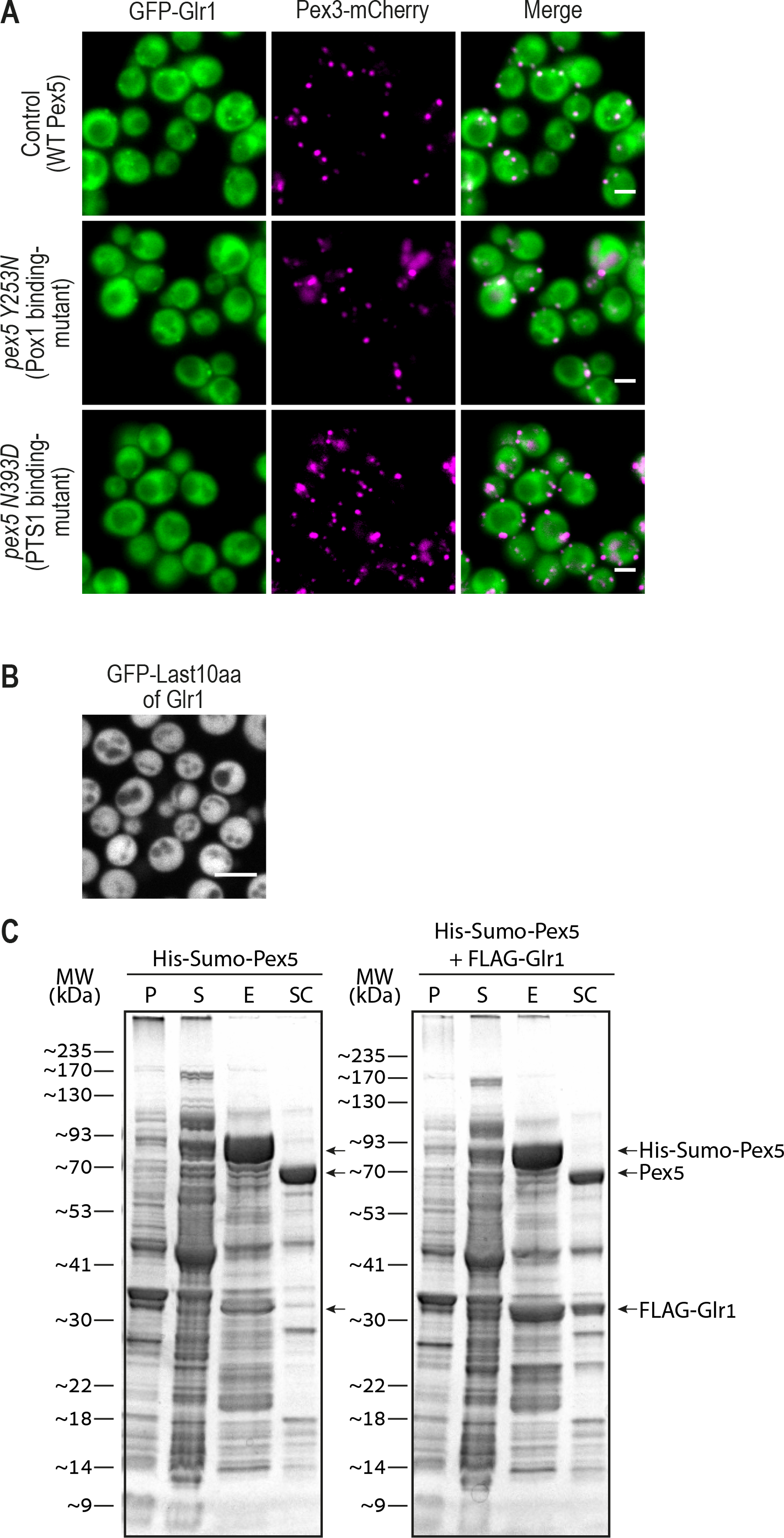
Glr1 is dependent on the PTS1 targeting route and binds Pex5 without containing a C’ PTS1 motif. (A) GFP-Glr1 peroxisomal localization was abolished upon point mutation in the PTS1-binding domain of Pex5 (Pex5N393D) (B) A strain expressing the last 10 amino acids of Glr1 fused to GFP at its C’ shows that they are not sufficient to target GFP to peroxisomes, hence Glr1 doesn’t contain a PTS1 motif (C) Direct and specific interaction between Glr1 and Pex5 was indicated by *in vitro* pull-down assays and SDS-PAGE analysis, showing Glr1 was co-eluted together with Pex5 following removal of the N’-His-Sumo tag on Pex5 (SC). P-Pellet fraction, S-Soluble fraction, E-Elution fraction, SC-Soluble fraction following bdSumo protease cleavage. For all micrographs, a single focal plane is shown. The scale bar is 5 um.

**Figure S6.**
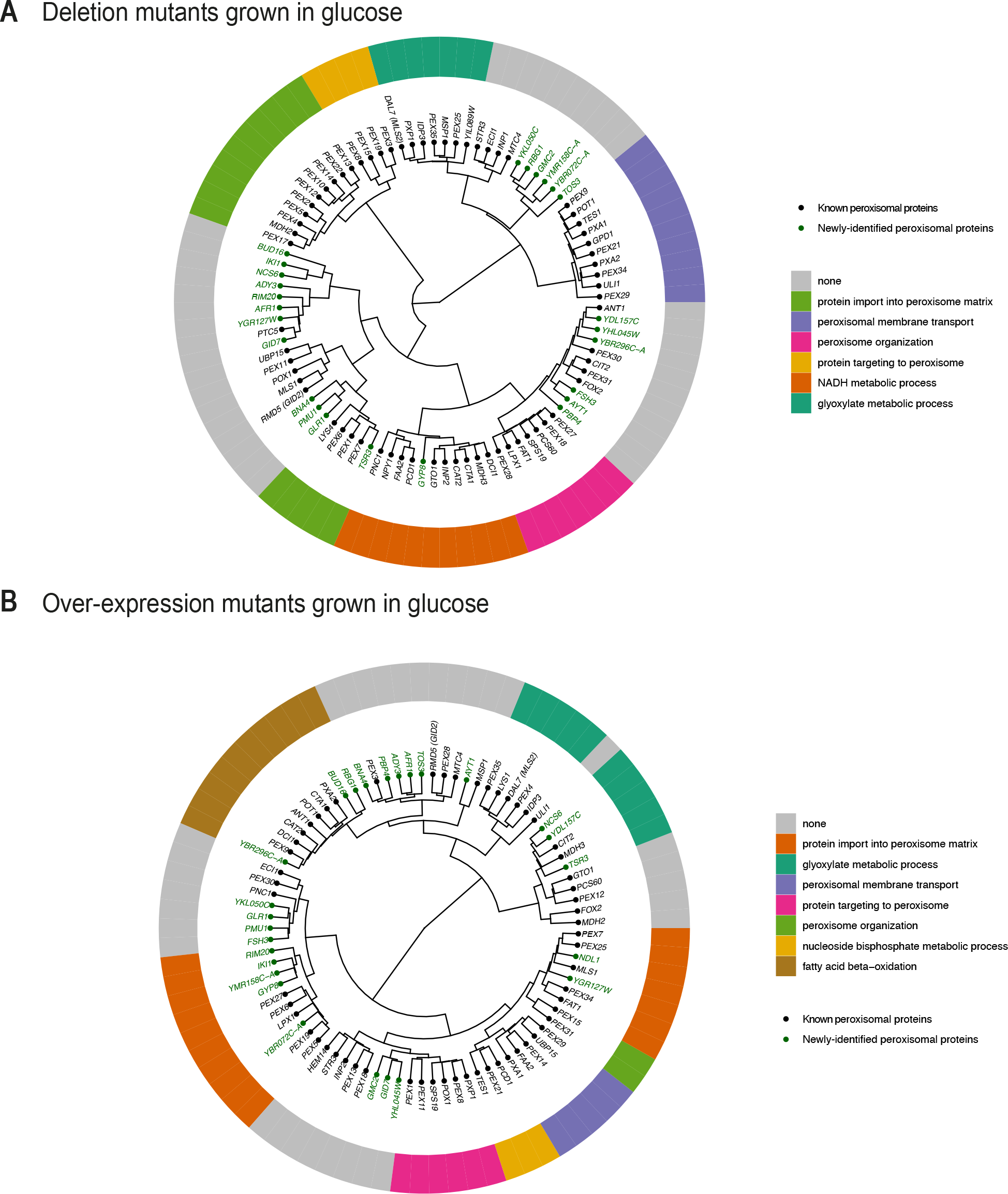
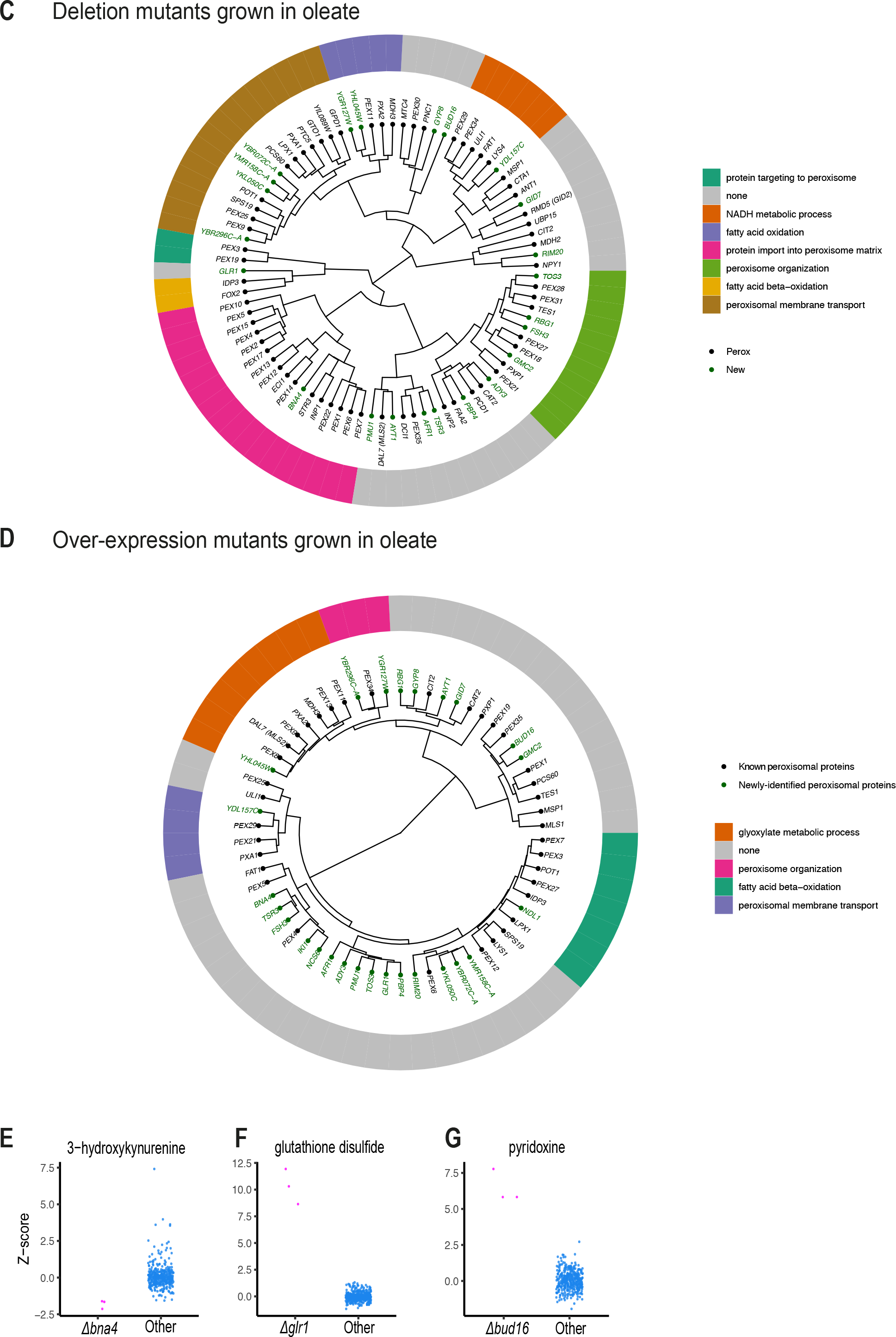
The hierarchical relationships between metabolomic fingerprints of different peroxisomal mutants. A circular dendrogram represents the hierarchal relationship between the metabolomic fingerprints of different peroxisomal mutants in each condition. (A) Deletion mutants grown in glucose (B) Over-expression mutants (*TEF2* promoter) grown in glucose (C) Deletion mutants grown in Oleate (D) Over-expression mutants (*TEF2* promoter) grown in Oleate. GO enrichment analysis of known peroxisomal genes in each cluster shows enrichment for the biological processes indicated. (E) The established activity of the Kynurenine 3-monooxygenase, Bna4, was detected by the metabolomic analysis of its deletion strain (reduction in 3-hydroxykynurenine). (F) The known activity of the glutathione oxidoreductase, Glr1, was captured by metabolomics, with elevation in glutathione disulfide molecules upon deletion of Glr1. (G) The putative role of Bud16 as a pyridoxal kinase was strengthened by the metabolomics data, showing *Δbud16* had elevated levels of pyridoxine.

**Figure S7.**
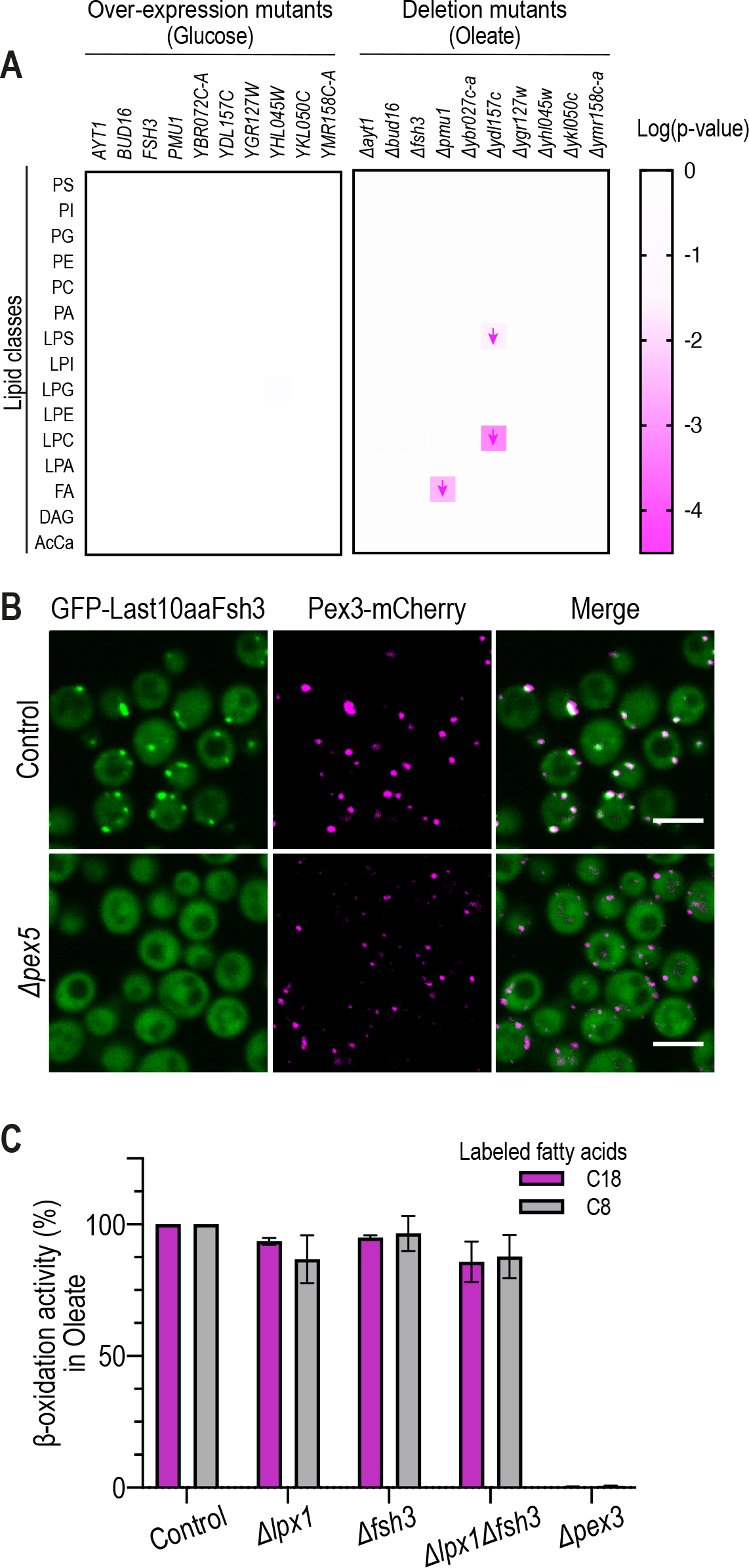
Functional assays for identifying mutants that affect lipid metabolism (A) Lipidomic analysis of peroxisomal deletion or over-expression mutants grown in glucose- or oleate-containing media was summarized in a heatmap. Arrows indicate directionality of fold-change (B) The last 10 amino acids of Fsh3 were sufficient to support the targeting of GFP to peroxisomes in a Pex5-dependent manner, demonstrating Fsh3 has a unique PTS1 motif. The scale bar is 5 µm. (C) A β-oxidation activity assay of *Δlpx1, Δfsh3*, and *Δlpx1Δfsh3* strains supplemented with labeled 8 carbon- or 18 carbon-fatty acids was measured following growth in oleate-containing media, showing no difference in β-oxidation activity between strains.

**Figure S8.**
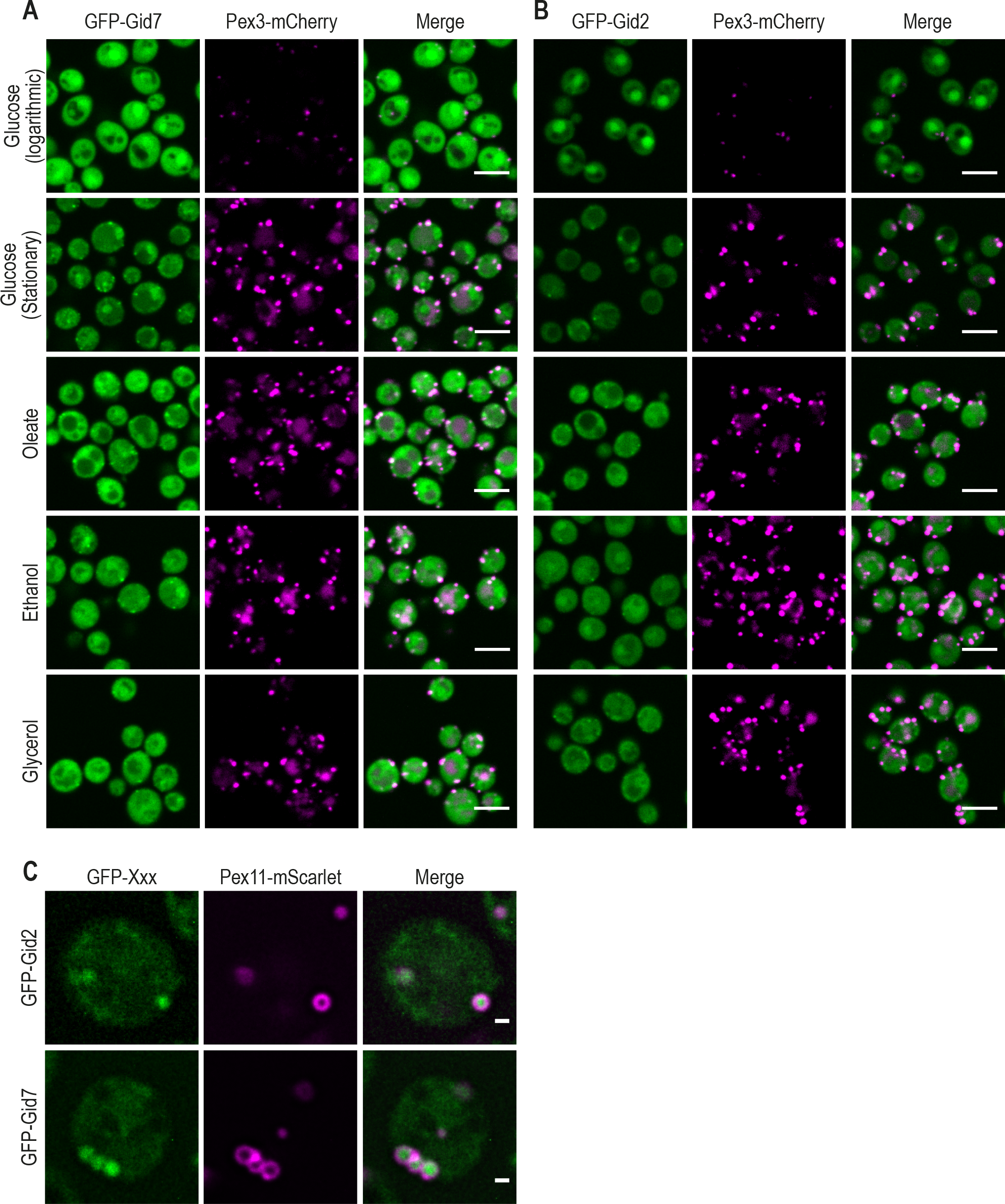
GID complex subunits Gid7 and Gid2 are targeted to the peroxisomal matrix only in gluconeogenic conditions. Micrographs of strains expressing (A) GFP-Gid7 and (B) GFP-Gid2 show that they localize to the cytosol when cells are grown in media supplemented with glucose, however, they partially localize to peroxisomes when cells are grown in media without glucose. Glucose (logarithmic) and Glucose (stationary) images for GFP-Gid7 are identical to main text figure 5C. The scale bar is 5 um. (C) High-resolution imaging shows both GFP-Gid2 and GFP-Gid7 localize to the peroxisomal matrix during growth in oleate. The scale bar is 500 nm. For all micrographs, a single focal plane is shown.

## Supplemental Information

Table S1. A list of known and newly-identified peroxisomal proteins and all summarized information curated for them in this study.

Table S2. Molecular dynamics analysis for predicting new PTS1 motifs among the newly-identified peroxisomal proteins.

Table S3. Average backbone H-bonds stability for peptide positions −1 and −3 used to detect likely binders within the Pex5 PTS1 binding cavity.

Table S4. Sample information and annotated ion intensities from the metabolomics profiling of all peroxisomal deletion mutants.

Table S5. Sample information and annotated ion intensities from the metabolomics profiling of all over-expressed (*TEF2* promoter) peroxisomal proteins.

Table S6. Sample information and annotated ion intensities from the metabolomics profiling of AYT1 deletion and over-expression (TEF2 promoter) mutants.

Table S7. Lipidomic analysis of ten newly-identified uncharacterized peroxisomal genes.

Table S8. Yeast strains and primers used in this study.

Table S9. Plasmids used in this study.

