## Supplemental Table 2 for "Systematic multi-level analysis of an organelle proteome reveals new peroxisomal functions"

**Table S2: Molecular dynamics analysis for predicting new PTS1 motifs among the newly-identified peroxisomal proteins.**

Percent of H-bond contacts (≤0.35nm) between backbone atoms of the peptide and Pex5 sidechains in the combined 180 ns trajectory (first row) and in the last 100 ns (second row).

| Peptide atom | O_t_ | | | | | N_-1_ | O_-2_ | | N_-2_ | O_-3_ | | N_-3_ | O_-5_ | | N_-5_ |
| --- | --- | --- | --- | --- | --- | --- | --- | --- | --- | --- | --- | --- | --- | --- | --- |
| Pex5 residue | Q359 | Asn393 | Asn495 | Arg496 | Arg526 | Asn393 | Asn495 | Asn530 | Asn530 | Asn503 | Tyr468 | Asn503 or Tyr468 | Tyr468 | Asn537 | Asn503 or Asn505  or Asn537 |
| hPex5*  YQSKL | - | 85.5  85.9 | 62.0  52.5 | 43.6  47.6 | 93.6  93.4 | 98.1  96.8 | 53.7  48.0 | 99.1  98.7 | 36.1  24.2 | 55.3  57.8 | - | 89.5  90.0 | - | 19.1  19.8 | 9.6  11.7 |
| Mls1  TDLSKL | 21.9  39.3 | 97.9  99.2 | 66.4  56.0 | 90.9  86.0 | 95.2  96.1 | 91.4  92.2 | 15.7  7.5 | 88.9  86.9 | 5.5  2.1 | 1.4  1.4 | 68.9  63.2 | 78.4  97.6 | 4.5  7.8 | 7.1  2.4 | 60.3  59.3 |
| Mls2  VDLSKL | 49.2  64.6 | 97.4  97.7 | 52.3  42.5 | 74.5  83.5 | 93.2  92.7 | 98.8  98.5 | 18.4  15.1 | 91.7  94.8 | 4.4  4.7 | 38.0  38.8 | 46.5  47.8 | 90.7  88.4 | 0.0  0.0 | 0.5  0.2 | 59.7  79.0 |
| Cit2  NIESKL | 23.5  42.3 | 96.9  95.9 | 82.2  83.1 | 69.8  81.7 | 96.1  95.7 | 96.9  95.9 | 37.6  38.0 | 77.7  96.1 | 13.0  5.9 | 13.6  18.4 | 71.9  81.2 | 61.9  65.0 | 0.1  0.2 | 10.3  1.6 | 35.6  33.2 |
| Pex8  SQSSKL | 0.1  0.2 | 90.8  93.1 | 59.8  54.2 | 81.8  83.9 | 89.1  90.8 | 86.3  81.1 | 36.1  25.1 | 80.0  87.8 | 13.9  6.1 | 45.0  53.4 | 28.5  31.8 | 65.3  74.9 | 3.0  0.0 | 1.4  2.3 | 45.5  38.4 |
| Str3  IKSSKL | 5.1  3.6 | 95.7  95.9 | 73.9  64.7 | 82.5  75.2 | 89.2  87.9 | 94.1  92.0 | 46.2  42.8 | 88.8  86.8 | 9.9  10.7 | 6.2  10.1 | 42.5  39.4 | 53.8  55.1 | 4.1  4.9 | 3.6  1.9 | 21.2  24.4 |
| Sps19  SMTSKL | 0.0  0.0 | 74.7  71.0 | 58.4  50.3 | 76.2  77.8 | 86.6  86.6 | 69.0  60.5 | 23.1  16.2 | 74.8  66.6 | 7.4  3.8 | 8.7  7.9 | 56.5  54.8 | 64.4  73.8 | 17.9  30.1 | 7.0  2.8 | 45.4  50.2 |
| Fox2  QAKSKL | 0.0  0.0 | 96.6  96.6 | 85.3  91.8 | 56.0  51.5 | 97.2  98.5 | 98.0  98.7 | 25.2  30.2 | 90.9  95.5 | 1.5  1.2 | 2.8  0.4 | 50.6  49.1 | 71.5  67.9 | 0.0  0.1 | 28.3  22.5 | 44.1  44.2 |
| Mdh3  LDSSKL | 6.2  11.2 | 91.1  92.0 | 64.6  55.8 | 90.5  93.0 | 92.2  89.7 | 91.6  87.2 | 30.0  25.1 | 73.4  67.2 | 14.6  10.5 | 0.1  0.2 | 61.4  59.5 | 46.9  58.6 | 1.8  3.2 | 5.6  7.8 | 50.5  56.1 |
| Pcs60  RNKSKL | 37.5  52.7 | 95.6  95.9 | 71.8  73.6 | 73.0  74.3 | 96.9  96.1 | 98.3  97.8 | 36.0  33.5 | 93.8  95.3 | 38.9  54.8 | 26.7  35.1 | 29.0  13.8 | 69.6  75.2 | 0.0  0.0 | 0.2  0.0 | 22.2  13.2 |
| Pxp2  SGVVKL | 18.3  24.7 | 97.7  97.6 | 53.9  61.8 | 82.2  81.8 | 91.5  93.0 | 96.7  96.9 | 11.6  14.4 | 67.8  77.9 | 3.6  6.2 | 8.4  8.5 | 44.8  28.5 | 23.8  7.5 | 1.2  0.4 | 3.1  3.4 | 22.4  22.8 |
| Idp3  KGMCKL | 9.0  16.2 | 76.9  75.0 | 35.3  34.2 | 64.0  60.4 | 82.6  87.2 | 68.2  66.2 | 27.2  31.0 | 50.6  50.4 | 22.5  32.2 | 1.8  0.0 | 48.4  59.2 | 75.6  76.2 | 21.6  21.3 | 0.9  0.2 | 37.7  36.3 |
| Aat2  TIEAKL | 9.8  17.6 | 97.9  98.8 | 76.5  76.1 | 85.5  87.6 | 97.5  97.6 | 98.1  98.8 | 49.2  47.0 | 91.9  94.4 | 11.8  14.4 | 14.6  17.1 | 64.9  66.3 | 34.0  35.8 | 0.0  0.0 | 14.1  0.0 | 56.4  52.2 |
| Cat2  KRKAKL | 0.0  0.0 | 93.3  92.5 | 72.4  67.8 | 76.2  85.2 | 91.0  92.0 | 93.9  93.2 | 39.0  37.8 | 77.6  74.0 | 2.4  2.1 | 12.6  5.3 | 73.7  78.2 | 75.2  78.7 | 8.9  12.6 | 3.4  2.5 | 45.9  36.2 |
| Faa2  VKTEKL | 3.1  4.0 | 90.9  97.9 | 77.2  78.7 | 77.2  92.7 | 88.1  96.5 | 98.1  98.8 | 44.4  48.1 | 77.4  94.0 | 1.2  2.2 | 22.2  10.1 | 62.9  47.8 | 59.2  47.2 | 0.7  0.2 | 0.1  0.2 | 10.4  7.8 |
| Lpx1  TTKQKL | 5.2  9.4 | 98.8  99.2 | 65.4  61.2 | 75.1  65.6 | 87.0  84.9 | 95.4  95.9 | 18.5  9.8 | 74.8  89.8 | 5.6  3.0 | 22.5  30.3 | 28.4  14.9 | 73.4  85.6 | 18.7  21.9 | 2.6  0.4 | 31.3  36.2 |
| Gto1  PDISRL | 48.1  48.7 | 95.3  93.6 | 40.6  38.5 | 77.9  74.6 | 90.3  90.3 | 97.2  97.1 | 11.6  7.6 | 76.5  76.1 | 7.0  4.4 | 11.3  11.5 | 62.9  81.3 | 72.9  74.6 | 0.2  0.3 | 0.7  0.9 | 45.6  56.8 |
| Lys1  KRSSRL | 0.0  0.0 | 97.6  96.9 | 62.0  48.3 | 81.3  87.4 | 89.3  83.6 | 91.6  86.7 | 32.3  18.2 | 55.5  38.2 | 7.4  0.2 | 6.9  10.0 | 83.8  92.1 | 68.4  79.6 | 1.1  0.3 | 0.6  0.2 | 34.7  35.4 |
| Eci1  QRKHRL | 0.1  0.0 | 79.5  79.6 | 35.5  31.0 | 85.1  84.2 | 69.6  57.7 | 68.3  71.1 | 15.3  10.6 | 65.0  73.9 | 6.2  7.5 | 0.0  0.0 | 12.3  8.7 | 99.0  98.6 | 29.9  32.0 | 0.2  0.3 | 20.1  20.6 |
| Pxp1  QNKPRL | 42.5  49.3 | 58.4  55.3 | 50.3  47.2 | 79.0  81.7 | 86.8  89.8 | 53.0  55.9 | 31.1  19.8 | 86.4  88.9 | 41.2  40.2 | 3.3  3.6 | 58.9  73.9 | -  - | 0.2  0.4 | 29.8  38.2 | 51.1  58.5 |
| Npy1  TSSSHL | 0.0  0.0 | 70.9  57.0 | 50.0  52.0 | 63.8  65.7 | 78.7  80.7 | 54.7  49.3 | 52.4  56.2 | 90.3  93.4 | 43.6  48.1 | 5.4  6.1 | 33.4  38.0 | 20.4  24.0 | 1.3  2.3 | 3.2  1.9 | 30.5  25.3 |
| Cta1  SSNSKF | 25.6  44.3 | 88.2  84.7 | 57.6  47.9 | 68.2  71.1 | 74.4  64.2 | 58.5  65.6 | 42.8  29.9 | 91.0  93.7 | 19.9  13.2 | 6.2  10.2 | 21.5  13.6 | 43.5  49.1 | 7.8  11.5 | 0.2  0.1 | 9.9  0.3 |
| Tes1  DIRAKF | 3.5  6.2 | 91.0  93.2 | 76.1  78.6 | 77.7  82.1 | 87.8  89.5 | 93.9  96.8 | 61.5  64.0 | 94.6  94.7 | 33.9  39.7 | 33.9  35.0 | 2.1  3.4 | 43.1  42.3 | 0.0  0.0 | 24.0  20.9 | 38.2  37.4 |
| Pmu1  ADRGRL | 2.8  0.0 | 78.4  79.7 | 17.8  10.3 | 49.7  40.5 | 62.2  59.3 | 6.0  0.0 | 46.4  35.6 | 72.8  59.5 | 56.9  47.7 | 0.8  0.0 | 4.9  4.3 | 4.5  0.4 | 0.0  0.0 | 19.9  32.6 | 52.5  68.8 |
| Ygr127w  RFKFKL | 0.0  0.0 | 36.1  2.8 | 30.7  28.2 | 54.8  39.0 | 77.0  74.8 | 33.2  2.0 | 17.3  17.8 | 45.6  42.0 | 0.0  5.2 | 0.0  0.0 | 28.6  32.1 | 97.4  96.2 | 57.5  46.1 | 0.0  0.0 | 22.7  15.4 |
| Bud16  YIYARL | 34.2  44.0 | 85.0  83.8 | 55.8  51.7 | 85.6  80.9 | 96.2  98.1 | 85.6  80.5 | 51.8  48.5 | 90.8  90.8 | 52.8  52.9 | 36.2  31.4 | 13.8  17.9 | 37.5  33.1 | 13.4  8.8 | 49.5  49.8 | 76.5  76.8 |
| Fsh3  DSLGKL | 1.2  2.2 | 88.6  80.8 | 47.4  36.9 | 85.8  83.7 | 79.4  77.5 | 86.5  81.2 | 47.8  41.8 | 65.3  51.5 | 29.8  39.8 | 5.2  2.3 | 27.0  13.5 | 13.7  7.9 | 2.0  2.6 | 7.7  1.2 | 15.5  2.1 |
| Mps1  FADYKI | 0.7  1.2 | 47.4  26.5 | 16.3  12.1 | 72.9  67.0 | 62.3  57.2 | 36.9  20.4 | 12.1  15.9 | 47.0  45.2 | 17.0  28.5 | 4.5  4.4 | 48.0  49.3 | 17.1  9.2 | 8.5  12.6 | 3.5  0.1 | 50.9  53.1 |
| Tos3  MSLYKL | 0.0  0.0 | 91.3  89.3 | 56.1  50.9 | 92.6  94.2 | 75.2  65.7 | 84.7  79.5 | 29.8  26.9 | 75.2  70.2 | 37.9  31.6 | 1.8  0.6 | 30.1  24.2 | 49.4  5.2 | 14.4  19.1 | 3.8  1.1 | 8.4  0.1 |
| Yhl045w  LGVAYL | 0.0  0.0 | 97.8  97.9 | 78.5  73.7 | 80.7  83.6 | 91.3  93.6 | 88.1  84.8 | 21.7  5.8 | 74.2  65.2 | 13.6  3.0 | 3.0  2.3 | 51.4  48.7 | 45.4  45.5 | 0.3  0.1 | 6.4  3.5 | 17.1  15.8 |
| Nud1  PTATNL | 30.1  14.7 | 73.6  59.2 | 41.4  38.0 | 45.4  34.3 | 66.9  72.6 | 64.6  48.1 | 22.2  32.2 | 45.6  42.5 | 36.6  36.4 | 5.8  4.8 | 37.8  27.2 | 0.6  0.5 | 0.6  0.1 | 0.0  0.0 | 0.0  0.0 |
| Gid7  WKISRN | 11.7  21.1 | 78.6  66.3 | 53.9  56.4 | 69.2  70.8 | 78.2  74.0 | 57.6  42.7 | 23.8  16.9 | 77.1  71.1 | 2.0  0.1 | 2.5  1.4 | 19.0  2.4 | 47.9  32.0 | 0.0  0.0 | 15.5  14.9 | 28.9  31.6 |
| Ybr072c-a  DSFVKT | 0.0  0.0 | 95.2  96.1 | 52.0  48.8 | 71.0  69.3 | 73.8  66.4 | 62.6  57.6 | 12.7  9.1 | 77.5  73.4 | 0.0  0.0 | 48.3  45.4 | 0.5  0.9 | 71.0  71.2 | 0.0  0.0 | 27.9  27.2 | 11.9  12.4 |
| Ncs6  LEKLSF | 2.3  4.1 | 58.5  49.2 | 80.6  72.4 | 80.9  72.1 | 88.3  84.5 | 24.0  10.5 | 67.2  57.8 | 93.6  92.2 | 28.9  21.8 | 0.1  0.2 | 12.4  8.6 | 24.8  21.8 | 18.1  13.4 | 12.4  16.6 | 34.6  46.1 |
| Ndl1  ATTSSV | 51.6  65.5 | 81.2  79.8 | 23.9  13.6 | 96.6  97.6 | 73.6  73.3 | 5.4  6.9 | 14.1  0.1 | 18.6  2.8 | 9.2  0.1 | 0.0  0.0 | 17.4  6.1 | 34.2  24.8 | 10.4  4.8 | 0.5  0.0 | 6.6  4.6 |
| Mdh2  SRSASS | 0.0  0.0 | 14.0  16.3 | 11.4  6.2 | 40.5  35.9 | 72.8  72.9 | 0.1  0.1 | 35.1  24.0 | 92.6  91.8 | 35.1  31.7 | 4.7  7.5 | 15.4  9.4 | 24.8  9.1 | 0.2  0.0 | 0.7  0.8 | 46.2  34.2 |

*)	Residues E344, N378, N462, N489, K490, N497, N499, R520, N524 and N531 of human Pex5 correspond to Q359, N393, Y468, N495, R496, N503, N505, R526, N530 and N537 of yeast Pex5.
