## Supplemental Table 3 for "Systematic multi-level analysis of an organelle proteome reveals new peroxisomal functions"

**Table S3: Average backbone H-bonds stability for peptide positions -1 and -3 used to detect likely binders within the Pex5 PTS1 binding cavity.**

| Protein | Position -1  Average and  Range (% of trajectory) | Position -3  Average and  Range (% of trajectory) |
| --- | --- | --- |
| 22 known  yPex5 cargos | 80.7±8.8  91.1 – 63.6 | 59.2±15.6  79.6 – 29.6 |
| Pmu1  ADRGRL | 42.8 | 5.1 |
| Ygr127W  RFKFKL | 46.4 | 62.9 |
| Bud16  YIYARL | 81.6 | 43.8 |
| Fsh3  DSLGKL | 77.5 | 23.0 |
| Mps1  FADYKI | 47.2 | 34.8 |
| Tos3  MSLYKL | 80.0 | 40.7 |
| Yhl045W  LGVAYL | 87.3 | 49.9 |
| Nud1  PTATNL | 58.4 | 22.1 |
| Gid7  WKISRN | 67.0 | 34.7 |
| Ybr072C-A  DSFVKT | 70.9 | 59.9 |
| Ncs6  LEKLSF | 66.5 | 18.7 |
| Ndl1  ATTSSV | 56.1 | 25.8 |
| Mdh2  SRSASS | 27.8 | 22.5 |
